# Targeting Dengue Virus NS3 Helicase: Biochemical and Computational Evaluation of Catechins from *Camellia sinensis* as Potential Therapeutic Leads^†^

**DOI:** 10.64898/2026.06.22.733882

**Authors:** Milla K. Wojciechowski, Luis Daniel Goyzueta-Mamani, Miguel Angel Chávez-Fumagalli, Edward L. D’Antonio

## Abstract

Dengue virus serotype 2 is a human pathogenic flavivirus encoding non-structural protein 3 (DEN2-NS3), a superfamily-2 viral helicase. DEN2-NS3 contains an N-terminal protease domain and a C-terminal RNA helicase/nucleoside 5’-triphosphatase (NTPase) domain essential for replication. The enzyme utilizes energy from NTP hydrolysis to translocate 3’-to-5’ along a duplex RNA substrate. Prompted by the high potency of (−)-epigallocatechin gallate (EGCG) against Zika virus, this study investigated if three catechins, EGCG, (−)-epicatechin gallate (ECG), and (−)-epigallocatechin (EGC), would be potent inhibitors of DEN2-NS3. Enzyme-inhibition assays demonstrated that the catalytic domain, DEN2-NS3(S171-K618), was strongly inhibited by the galloylated catechins (EGCG and ECG), verified by an enzyme-coupled confirmatory assay. Inhibition constants (K_i_) and suggestive inhibition modes relative to NTPase activity were K_i_ = 400 ± 86.6 nM for EGCG (mixed-mode), K_i_ = 550 ± 250 nM for ECG (uncompetitive), and K_i_ = 18.3 ± 4.2 µM for EGC (mixed-mode). The coronavirus inhibitor SSYA10-001 inhibited DEN2-NS3 (K_i_ = 10.2 ± 0.3 µM, mixed-mode). Computational workflows using SiteMap identified a predicted druggable pocket associated with the RNA-binding region, involving residues ASP290, ARG387, ASP409, MET429, HIS487, ASP541, ARG599, and ASP603. Catechins were analyzed through 200-ns molecular dynamics simulations to evaluate binding stability. Computational results revealed that EGCG and ECG maintained high stability, forming highly persistent shared amino acid contacts (>45% occupancy) with ASP603, ARG599, ASP541, and ARG387. Biochemical and computational data support these galloylated catechin leads, suggesting a plausible binding model within an amphipathic pocket. Future structural optimization into stable prodrug derivatives could yield promising antiviral candidates.

**Highlights:**

- EGCG and ECG are highly potent inhibitors of the DEN2-NS3 helicase.
- Computational modeling suggests that galloylated catechins can be accommodated within a predicted pocket associated with the RNA-binding region.
- MD simulations (200-ns trajectories) revealed continuous binding stability.
- Key interaction residues include ASP603, ARG599, ASP541, and ARG387.
- Experimental enzyme-inhibition corroborated with computational findings.

**Graphical Abstract:** 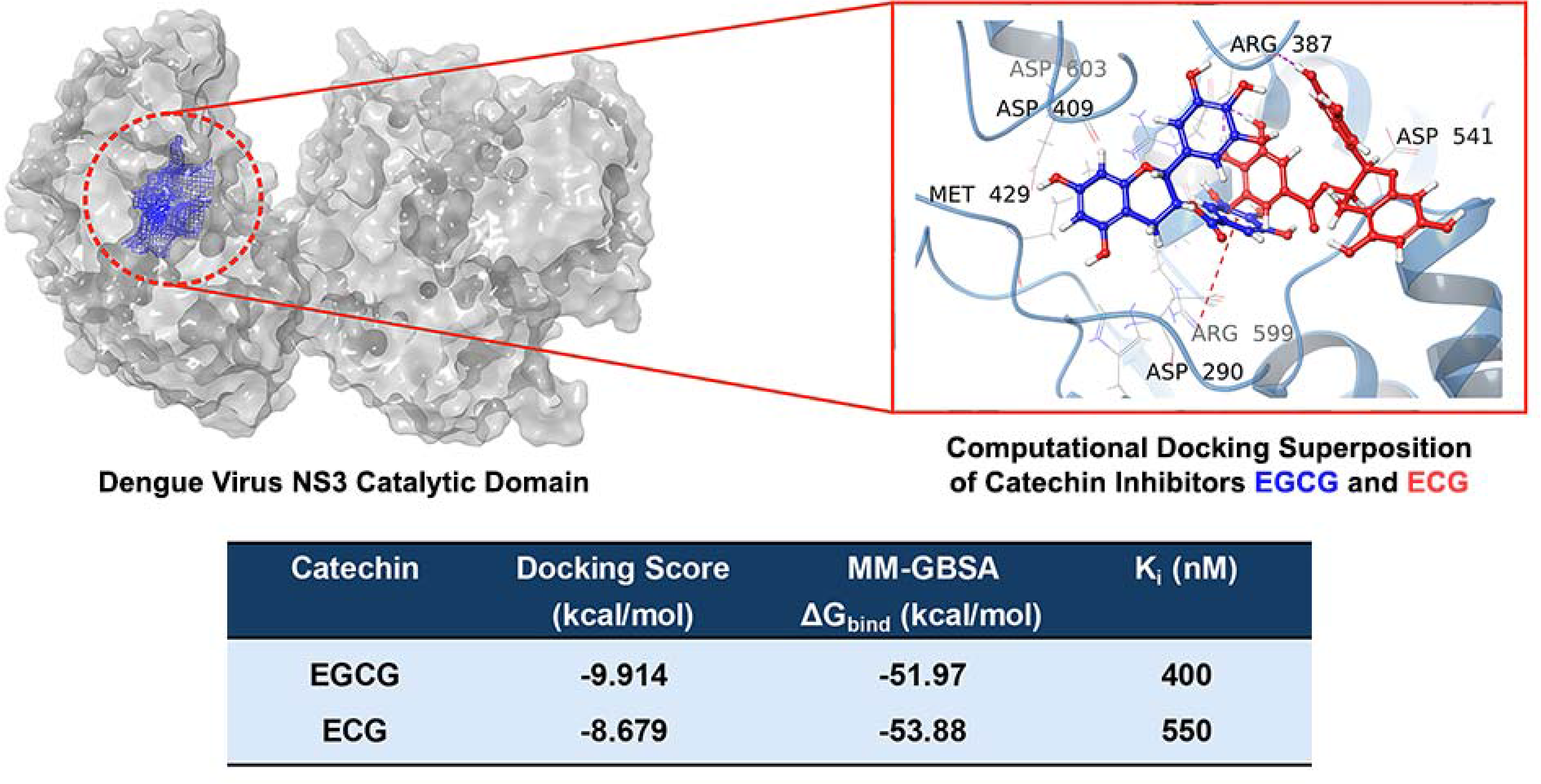

## 1. Introduction

Dengue virus (DENV) is a mosquito-transmitted pathogen that poses a persistent global health challenge, with an estimated 100 – 400 million infections occurring each year, of which approximately 96 million result in symptomatic illness [1]. Transmitted primarily by *Aedes aegypti* mosquitoes, DENV is endemic in over 100 countries, particularly in tropical and subtropical regions where favorable environmental conditions support year-round transmission [2]. Although vaccine availability remains limited, it is gaining traction. There are only two main vaccines that have been distributed globally, which include Dengvaxia (Sanofi Pasteur) [3] and Qdenga (TAK-003) (Takeda) [4]. A third emerging vaccine in the developmental stage is the single-dose Butantan-Dengue Vaccine (Instituto Butantan) [5]. There are no specific antiviral therapies approved for Dengue virus infection [1]. Among the four distinct serotypes of the virus (DENV-1 to DENV-4), secondary infection by a heterologous serotype increases the risk of developing severe disease forms such as Dengue hemorrhagic fever or Dengue shock syndrome through antibody-dependent enhancement [6]. Consequently, discovering small-molecule inhibitors targeting essential enzymes from the virus has become a critical area of research. One promising target is the Dengue virus non-structural protein 3 (DENV-NS3), which plays an essential role in viral RNA replication and is highly conserved among flaviviruses [7]. Natural polyphenolic compounds, particularly some catechins derived from *Camellia sinensis* (the tea plant) (**Figure 1**), have been shown to exhibit antiviral [7] and anticancer [8, 9] properties (e.g., (−)-epigallocatechin gallate (EGCG)). Furthermore, EGCG was found to have antiviral activity against DENV in Vero cells with an IC_50_ of 18.0 ± 1.0 µM [10], and Kumar and colleagues showed that EGCG was a potent inhibitor of NS3 helicase/NTPase from Zika virus (ZIKV) [11]. These findings highlight the potential of repurposing green tea catechins as compounds of interest targeting DENV-NS3, for which it is still unclear whether these compounds can act as inhibitors.

**Figure 1:**
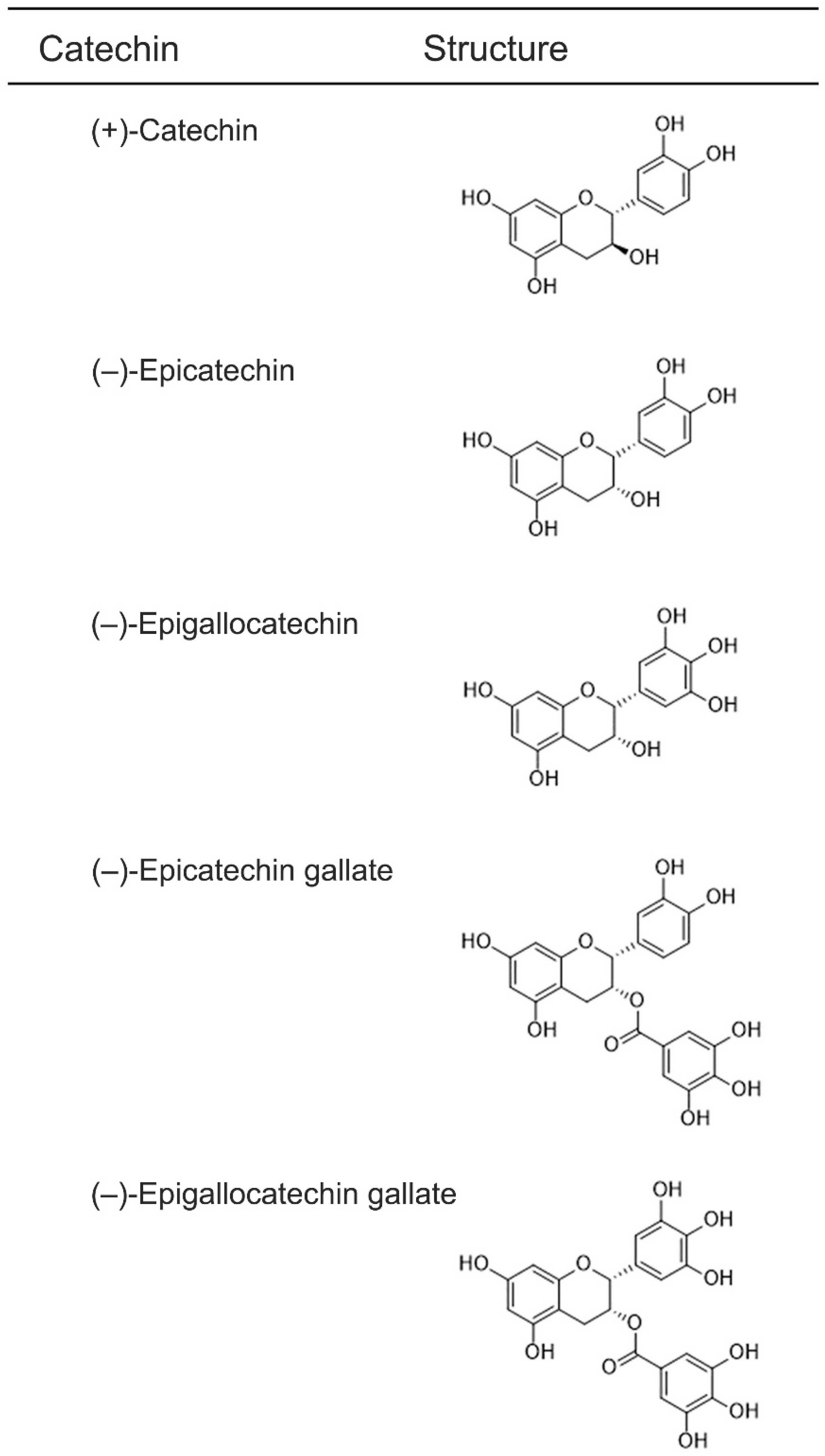
Chemical structures of naturally produced catechins from *Camellia sinensis*. Catechin compounds, shown from top to bottom, include (+)-catechin, (−)-epicatechin (EC), (−)-epigallocatechin (EGC), (−)-epicatechin gallate (ECG), and (−)-epigallocatechin gallate (EGCG).

DENV-NS3 is a key multifunctional enzyme required for viral replication. It contains two domains, such as an N-terminal serine protease domain and a C-terminal RNA helicase/nucleoside 5’-triphosphatase (NTPase) domain [7]. The protease domain associates with NS2B in order to cleave the viral polyprotein into individual non-structural proteins required for replication. These seven proteins include NS1, NS2A, NS2B, NS3, NS4A, NS4B, and NS5. In addition to its importance in polyprotein processing, DENV-NS3 also interacts directly with NS4B [7,12,13]. The helicase/NTPase domains use energy from NTP hydrolysis to unwind RNA substrates and help with viral RNA synthesis [14,15]. As a superfamily-2 (SF2) viral helicase [11], DENV-NS3 contains several conserved motifs that coordinate ATP binding with its hydrolysis, RNA binding, and this allows the enzyme to translocate in a 3′-to-5′ direction while separating duplex RNA strands (Eq. 1a, **Figure 2a**). DENV-NS3 can also unwind dsDNA, but it has a higher preference for dsRNA [14,16]. A short polypeptide segment linking the protease and helicase domains permits the two domains to move relative to each other in a flexible manner, which helps the protease to process the viral polyprotein while the helicase unwinds RNA efficiently [10,13,17]. The protease, helicase, and NTPase activities are essential for viral replication, and NS3 is consequently an important target for antiviral therapeutic drug development [10,14].

**Figure 2:**
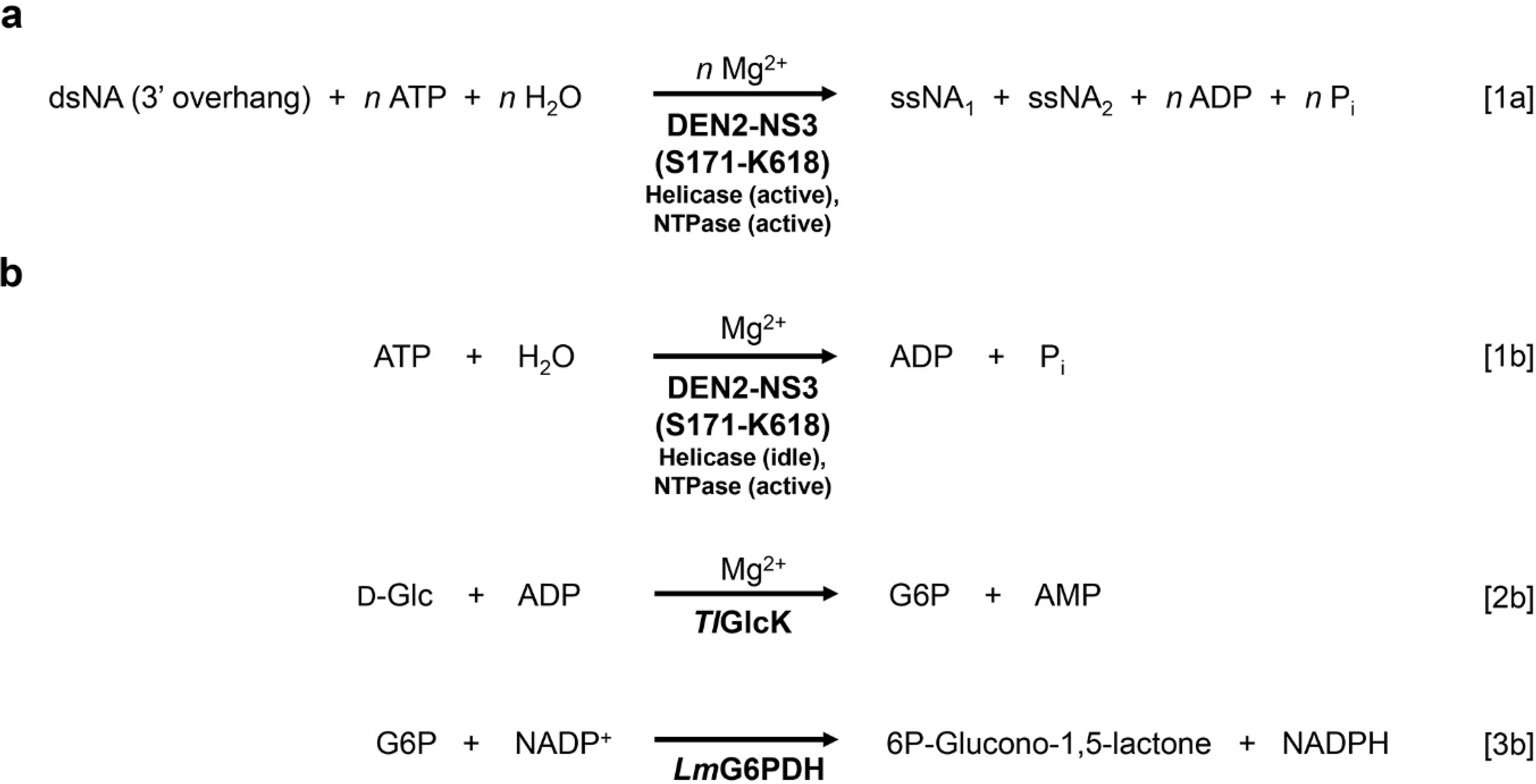
Reactions and fluorometric assay mechanism of the DEN2-NS3(S171-K618) helicase/nucleoside 5’-triphosphatase (NTPase). (a) The multifunctional enzyme DEN2-NS3(S171-K618) helicase/NTPase functions to catalyze the unwinding of double-stranded nucleic acid (dsNA). ATP, H_2_O, Mg^2+^, and the substrate dsNA (preferably dsRNA bearing a 3’-ended overhang), undergo conversion into products consisting of single strands of nucleic acid (ssNA), ADP, and inorganic phosphate. (b) DEN2-NS3(S171-K618) helicase/NTPase enzyme-coupled fluorometric-based assay, as performed in this study. The assay functions with the helicase domain in the idle state and the NTPase domain in the active state. Through the coupling of the ADP-dependent *Thermococcus litoralis* glucokinase (*Tl*GlcK) and *Leuconostoc mesenteroides* glucose-6-phosphate dehydrogenase (*Lm*G6PDH) via the progression of ADP through the assay, NADPH becomes the final product that is fluorometrically active (λ_ex_ = 340 nm, λ_em_ = 485 nm).

Prior research by Coronado et al. demonstrated that DENV-NS3 could be moderately inhibited by green tea catechins, such as EGCG and (−)-epigallocatechin (EGC). Their study revealed that inhibitory activity was observed from an assay of the protease domain (DENV-NS3_pro_), but not from the helicase/NTPase domain (DENV-NS3_hel_). IC_50_ values were measured as 6.3 µM for EGCG and 145 µM for EGC, suggesting that these catechins were not very potent small-molecule inhibitors of DENV-NS3_pro_ [18]. On the other hand, Kumar et al. demonstrated that EGCG was a potent inhibitor of Zika virus non-structural protein 3 (ZIKV-NS3) (an SF2 viral helicase) that exhibited an IC_50_ value of 295.7 nM [11]. Through structural comparison of the full-length protein sequence of ZIKV-NS3 (GenBank accession no. ASK51714.1, full polypeptide) vs. DENV-NS3 (GenBank accession no. WVS18090.1, position 1476 – 2093) by BLASTp, their protein sequence identity is 66.7%. This level of sequence identity supports the hypothesis that EGCG and related catechins may also inhibit the DENV-NS3 helicase domain. Since the DENV protease domain was probed for inhibition in the past, but not the helicase domain, we were motivated by Kumar et al. to observe for potent inhibition of DENV-NS3_hel_ by EGCG, and possibly other green tea catechins. Furthermore, our laboratory has recently developed a novel assay for the nsp13 helicase/NTPase in Severe Acute Respiratory Syndrome Coronavirus 2 (SARS-CoV-2), a superfamily-1B (SF1B) viral helicase, which utilizes an enzyme-coupled reaction scheme [19]. Briefly, this helicase/NTPase assay combines three enzyme-coupled steps involving the ADP-dependent *Thermococcus litoralis* glucokinase (*Tl*GlcK), *Leuconostoc mesenteroides* glucose-6-phosphate dehydrogenase (*Lm*G6PDH), and *Clostridium kluyveri* diaphorase (*Ck*DIA). The overall reaction produces a water-soluble tetrazolium chromophore called iodonitrotetrazolium formazan that can be helpful for rapid colorimetric assay determinations. However, using the SARS-CoV-2 helicase/NTPase assay as a starting point, we made key changes to probe only the NTPase activity. Moreover, we studied the DENV serotype 2 (henceforth referred to as DEN2) NS3 protein, truncating the N-terminal protease domain to separate the C-terminal helicase/NTPase domain, DEN2-NS3(S171-K618). The NTPase activity was held in an active state, but the helicase activity was held in an idle state (Eq. 1b, **Figure 2b**). This modification was accomplished by not including its preferred substrate, 3’-ended overhang double-stranded nucleic acid (dsNA), in the reaction mixture of the assay. The enzyme-coupled reaction involved two enzymes, such as *Tl*GlcK (Eq. 2b, **Figure 2b**) and *Lm*G6PDH (Eq. 3b, **Figure 2b**). With *Lm*G6PDH as the final enzymatic step, it produces the fluorophore NADPH (λ_ex_ = 340 nm, λ_em_ = 485 nm) [20], and the assay has a 1:1 stoichiometric relationship for the moles of ATP consumed to the moles of NADPH produced.

In the present work, we investigated three DENV-NS3_hel_ inhibitors that were suggestive of being either mixed-mode or uncompetitive relative to the NTP-binding site, and these compounds were catechins derived from the tea plant. The galloylated catechins EGCG and ECG revealed a very high inhibition towards DENV-NS3_hel_ with nanomolar inhibitory constants. As such, these compounds were prioritized for a computational workflow involving molecular docking and 200-ns MD simulations.

## 2. Materials and Methods

### 2.1. Chemicals and Reagents

20-mer ssDNA (5’-AGAGTTTGATCCTGGCTCAG-3’) and 5’-overhang 31/18-mer duplex dsDNA (utilized as a putative effector for NTPase activity from DEN2-NS3(S171-K618)) were purchased from Integrated DNA Technologies (Coralville, IA). The oligonucleotide sequences for the duplex were as follows: 31-mer ssDNA (5’-to-3’): CGCAGTCTTCTCCTGGTGCTCGAACAGTGAC; and 18-mer ssDNA sequence (5’-to-3’): GTCACTGTTCGAGCACCA). Isopropyl β-D-thiogalactopyranoside was purchased from Biosynth (San Diego, CA). (−)-Epigallocatechin gallate (EGCG) was purchased from Enzo Life Sciences (Farmingdale, NY). 3-[(2-Nitrophenyl)sulfanylmethyl]-4-prop-2-enyl-1*H*-1,2,4-triazole-5-thione (SSYA10-001) was purchased from MolPort (Beacon, NY). Ethylenediaminetetraacetic acid tetrasodium salt hydrate, (−)-epicatechin gallate (ECG), (−)-epigallocatechin (EGC), imidazole, bovine pancreas deoxyribonuclease I (DNase I), bovine pancreas ribonuclease A (RNase A), triethanolamine (TEA), β-nicotinamide adenine dinucleotide phosphate hydrate (NADP^+^), adenosine 5’-triphosphate disodium salt hydrate (ATP, >99%), Terrific broth, kanamycin sulfate, and dimethyl sulfoxide (DMSO) were purchased from Sigma-Aldrich (St. Louis, MO). *Leuconostoc mesenteroides* glucose 6-phosphate dehydrogenase (*Lm*G6PDH; E.C. 1.1.1.49) was purchased from Worthington Biochemical Corporation (Lakewood, NJ). Luria-Bertani (LB) broth, lysozyme (type VI), cobalt-nitrolotriacetic acid resin, protease inhibitor tablets (EDTA-free), D-glucose, glycerol, magnesium chloride, sodium phosphate dibasic, potassium phosphate monobasic, DL-dithiothreitol, and all other chemicals were purchased from Fisher Scientific (Hampton, NH).

### 2.2. Cloning and Preparation of the cDNA for DEN2-NS3(S171-K618) and TlGlcK

Gene synthesis was performed for the gene of *Dengue Virus Type 2 (DEN2) Helicase NS3 catalytic domain (S171 – K618)* (note: the truncation was established based on the protein sequence of the full-length NS3 gene (GenBank accession number WVS18090.1; position 1476 – 2093)). The catalytic domain was cloned into a kanamycin–resistant pET-28a(+) *Escherichia coli* expression vector at restriction sites 5’ NcoI and 3’ HindIII at Azenta Life Sciences, Inc. (South Plainfield, NJ). The plasmid can be obtained through Addgene.org (Addgene plasmid no. 258837) (https://www.addgene.org/258837/). *Thermococcus litoralis* (DSM 5473) ADP-dependent glucokinase (GenBank accession number Q7M537.1) was previously cloned into a kanamycin-resistant pET-28a(+) *E. coli* expression vector [19], and can be obtained through Addgene.org (Addgene plasmid no. 258836) (https://www.addgene.org/258836/). Each construct encoded an N-terminal hexahistidine tag, through the segment MGRGSHHHHHHGMA, that preceded SER171 in DEN2-NS3(S171-K618) and the start methionine in *Tl*GlcK. Codons for both plasmids were optimized to be used in protein expression of *E. coli*. The plasmid constructs were designated as pET-*DEN2-NS3(S171-K618)* and pET-*Tl*GlcK, respectively.

### 2.3. Expression and Purification of Recombinant DEN2-NS3(S171-K618)

Transformation of the plasmid pET-*DEN2-NS3(S171-K618)* was performed by the heat shock method using the *E. coli* strain BL21(DE3) (New England Biolabs) and colonies were grown on LB-agar plates with a concentration of kanamycin at 50 µg/mL. Starter cultures containing 5 mL of LB broth, 50 µg/mL of kanamycin, and a single colony of *E. coli* from the transformation step were incubated at 37 °C and shaking at 220 rpm for 7 h using a New Brunswick Scientific C24 incubator shaker. The starter cultures were used to inoculate six 2-L culture flasks containing 500 mL of Terrific broth modified (Sigma-Aldrich) supplemented with 0.4% (v/v) glycerol and including kanamycin at a concentration of 50 µg/mL that followed a 15 h incubation at 37 °C and shaking at 220 rpm. The culture flasks used autoclaved cheesecloth for lids for the air flow. The 500 mL cultures were induced with isopropyl β-D-thiogalactopyranoside to a final concentration of 1.0 mM and were incubated for an additional 25 h at 25 °C with shaking at 220 rpm. The *E. coli* was centrifuged at 4,100 rpm using a Beckman Coulter centrifuge equipped with a swinging bucket rotor, and the resultant cell paste was recovered using a spatula (50.1 g) followed by storing at -80 °C for at least 12 h (overnight). The frozen *E. coli* cell paste was thawed out from -80 °C to room temperature followed by resuspension in lysis buffer [50 mM 4-(2-hydroxyethyl)piperazine-1-ethanesulfonic acid (HEPES) (pH 7.0), 300 mM NaCl]. Cell lysis was accomplished by adding lysozyme and the cell suspension was stirred for 1 h at 4 °C before adding EDTA-free protease inhibitor tablets, after which the cell lysate was sonicated for 30 min in a water-bath sonicator (FS20, Fisher Scientific) and stored at -20 °C overnight. The cell lysate was thawed to room temperature and DNase I and RNase A at final concentrations of 10 μg/mL and 16 μg/mL, respectively, were added and stirred at 4 °C for 1 h, followed by overnight freezing at -80 °C. The cell lysate was thawed to room temperature and centrifuged at 11,400 rpm in a Beckman-Coulter centrifuge with the F0850 fixed-angle rotor for 40 min at 4 °C and the resulting supernatant was recovered and loaded onto a Co^2+^-affinity chromatography column [1.5 cm (internal diameter) x 4.0 cm (bed height)] that was pre-equilibrated with mobile phase A [50 mM HEPES (pH 7.0), 300 mM NaCl]. Mobile phase A was used as an initial wash to aid in the elution of most protein impurities. An isocratic step was applied from 0 – 13% mobile phase B, where mobile phase B was 50 mM HEPES (pH 7.0), 300 mM NaCl, 150 mM imidazole. After the UV absorbance (λ = 280 nm) on the chromatogram reached baseline, an isocratic step was applied to elute at 100% mobile phase B. Fractions of DEN2-NS3(S171-K618) were then pooled and concentrated down to approx. 5 mL using an Amicon Ultra-15 (MWCO = 30 kDa) centrifugal concentrator (Millipore). Meanwhile, a 16/600 size-exclusion chromatography column (GE Life Sciences) was pre-equilibrated with three column volumes of mobile phase [50 mM TEA (pH 7.6), 150 mM NaCl]. The 5 mL sample was loaded onto the column using a superloop and the main fraction band was recovered. UV-visible spectrophotometry (Agilent 8453) was used to determine whether nucleic acids were present by monitoring the wavelength at λ = 260 nm. Since DEN2-NS3(S171-K618) had a spectrum with a peak at 280 nm, it was determined that no nucleic acids were bound to the enzyme or present in the solution. Sodium dodecyl sulfate – polyacrylamide gel electrophoresis revealed the final fractions to be approximately 98% pure, by visual inspection. The pure fractions were pooled and concentrated to 1.0 mg/mL (18.9 x 10^-6^ mol/L) [ε_280_= 66,140 M^-1^ cm^-1^ [21]; M.W. = 52,845 g/mol (monomer of His-tagged DEN2-NS3(S171-K618))].

### 2.4. Expression and Purification of Recombinant TlGlcK

The pET-*Tl*GlcK plasmid was transformed by the heat shock method using *E. coli* strain BL21(DE3), and the cell colonies were grown on LB-agar plates with 50 µg/mL of kanamycin. Starter cultures containing 5 mL of LB broth, 50 µg/mL of kanamycin, and a swabbed colony of *E. coli* (from the transformation step) were incubated at 37 °C with shaking at 250 rpm for 8 h using an incubator shaker. The starter cultures were used to inoculate six 2-L culture flasks containing 500 mL of Terrific broth modified (Sigma-Aldrich) supplemented with 0.4% (v/v) glycerol that included 50 µg/mL of kanamycin. The incubation time was set to 16 h and held at a temperature of 37 °C with shaking at 220 rpm. The culture flasks used lids that were autoclaved cheesecloth squares (127-mm x 127-mm) held on by rubber bands, for improved air flow. The 500-mL cultures were induced with isopropyl β-D-thiogalactopyranoside to a final concentration of 1.0 mM and were incubated for an additional 24 h at 25 °C with shaking at 220 rpm in an incubator shaker. The *E. coli* was centrifuged at 4,100 rpm, and the resultant cell paste was recovered and stored at -80 °C overnight. The frozen cell paste was thawed from -80 °C to room temperature and resuspended in lysis buffer [50 mM HEPES (pH 7.0), 150 mM NaCl]. Cell lysis was performed by adding lysozyme and the cell suspension was stirred for 1 h at 4 °C before adding EDTA-free protease inhibitor tablets. Then the cell lysate was sonicated for 30 min in a water-bath sonicator followed by being stored at -20 °C overnight. The cell lysate was thawed to room temperature and exposed to two nucleases (final concentrations: 10 μg/mL of DNase I and 16 μg/mL of RNase A) and stirred at 4 °C for 1 h, followed by overnight freezing at -80 °C. The cell lysate was thawed to room temperature and centrifuged at 11,400 rpm for 45 min at 4 °C, and the supernatant was recovered, followed by loading onto a Co^2+^-affinity chromatography column [1.5 cm (internal diameter) x 4.0 cm (bed height)] pre-equilibrated with mobile phase A [50 mM HEPES (pH 7.0), 300 mM NaCl]. Mobile phase A was used as an initial wash to aid in the elution of protein impurities, followed by an isocratic step to 13% mobile phase B [100% MP-B = 50 mM HEPES (pH 7.0), 300 mM NaCl, 150 mM imidazole]. After the UV absorbance (λ = 280 nm) on the chromatogram reached baseline, a gradient elution from 13 – 100% mobile phase B (50 mL gradient) was implemented to elute most of the other impurities from the column. Fractions of *Tl*GlcK resulting from the Co^2+^-affinity chromatography step were pooled and concentrated to 5 mL using an Amicon Ultra-15 centrifugal concentrator (MWCO = 30 kDa). The *Tl*GlcK sample was subsequently loaded onto a 16/600 size-exclusion column pre-equilibrated with filtered mobile phase [50 mM TEA (pH 7.6), 150 mM NaCl]. Sodium dodecyl sulfate – polyacrylamide gel electrophoresis revealed the final fractions to be >99% pure. The pure fractions were pooled and concentrated to 1.0 mg/mL [ε_280_ = 50,310 M^-1^ cm^-1^ [21]; M.W. = 55,192 g/mol (monomer of His-tagged *Tl*GlcK)].

### 2.5. Kinetics and Inhibition Studies of DEN2-NS3(S171-K618) by Catechin Compounds

The NTPase binding site was probed for enzyme inhibition. The determination of inhibition constant (K_i_) values for inhibitors targeting DEN2-NS3(S171-K618) was performed by a three-enzyme fluorometric assay (**Figure 2b**). The assay was based on the formation of NADPH (fluorophore) in aqueous buffered solution at pH 7.6 (λ_ex_= 340 nm, λ_em_= 485 nm) [20]. DEN2-NS3(S171-K618) in the presence of a putative effector, such as a 5’-ended overhang 31/18-mer dsDNA, along with ATP, Mg^2+^, and H_2_O causes the formation of ADP and inorganic phosphate (P_i_). *Tl*GlcK in the second step combines with ADP and D-glucose to form G6P and AMP. Finally, *Lm*G6PDH combines with G6P and NADP^+^ to form 6-phospho-glucono-1,5-lactone and NADPH. DEN2-NS3(S171-K618) was first tested for enzyme activity in the absence of inhibitor. For the time optimum experiment, the concentration of 5’-ended overhang 31/18-mer dsDNA was held constant at a final assay concentration (F.A.C.) of 17.9 µM, but from a re-quantification of the stock then gave an actual concentration of 20.0 µM. ATP concentrations ranged from 0.0156 – 2.000 mM. F.A.C.s for the other reagents in the assay were as follows: 1.75 mM D-glucose, 0.477 mM NADP^+^, 6.95 mM MgCl_2_, 5.6 µg/mL *Tl*GlcK, 5.6 µg/mL *Lm*G6PDH, and 5.6 µg/mL DEN2-NS3(S171-K618). The reaction was carried out in a buffered solution [50 mM TEA (pH 7.6), 150 mM NaCl], and the reaction mixtures were added to a black, round-bottom microplate. Three negative controls were also included, which had ATP and 31/18-mer duplex substituted for the assay buffer, thus not having any ATP or duplex in the reaction. Reactions were performed at room temperature (25 °C), initiated with DEN2-NS3(S171-K618), utilized a reaction time of 200 s, and were immediately scanned in a VANTAstar-F multi-mode microplate reader (BMG LABTECH, Inc.; Cary, NC, USA) without any reaction-stopping solution (thus, reactions were not chemically terminated). The reaction time of 200 s was determined from a time optimum evaluation (**Figure 3**), and the reaction (which was initiated by the addition of ATP) consisted of the following reagents at their F.A.C. values: 17.9 µM 5’-ended overhang 31/18-mer dsDNA, 1.78 mM ATP, 1.75 mM D-glucose, 0.477 mM NADP^+^, 6.95 mM MgCl_2_, 6.0 µg/mL *Tl*GlcK, 6.0 µg/mL *Lm*G6PDH, and 6.0 µg/mL DEN2-NS3(S171-K618). The reaction was performed in a buffered solution [50 mM TEA (pH 7.6), 150 mM NaCl] at 22 °C with a sample volume of 168 µL. A negative control test was also performed, in which 5’-ended overhang 31/18-mer dsDNA and DEN2-NS3(S171-K618) were not added into the reaction mixture, but their volumes were replaced with assay buffer. The sample volume was also 168 µL. The negative control showed very minimal activity, as represented by an absorbance of 0.06 that was generated at λ = 340 nm by 322 s. Fluorescence measurements of NTPase activity corresponded to NADPH relative fluorescence (λ_ex_ = 340 nm, λ_em_ = 485 nm) that correlated to a 1:1 stoichiometry of ADP formation. All measurements were performed in triplicate, and the Michaelis-Menten parameters were determined, such as K_M_, k_cat_, k_cat_/K_M_, and V_MAX_.

**Figure 3:**
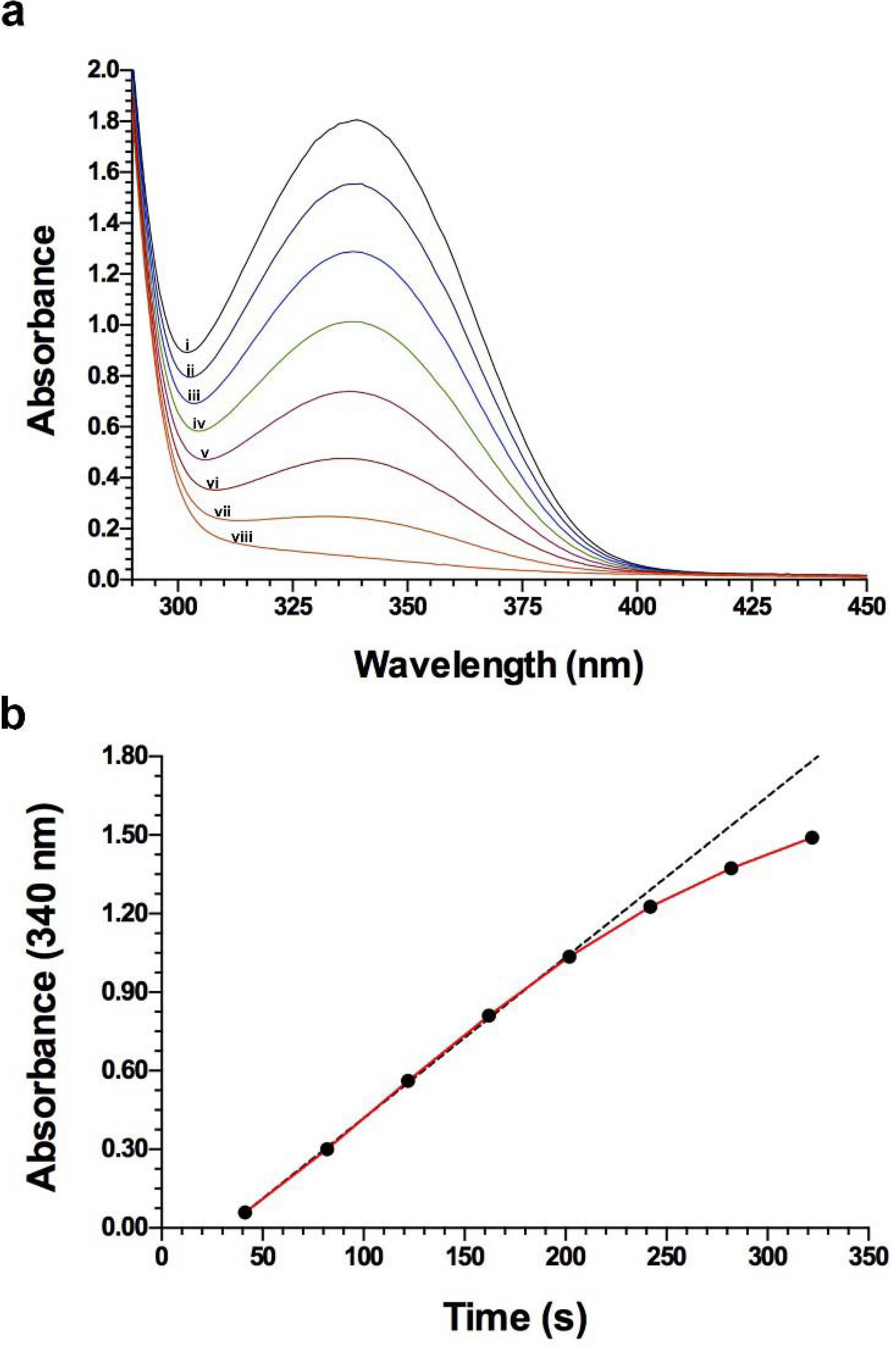
Time optimum determination of the DEN2-NS3(S171-K618) NTPase activity assay. (a) The UV–visible absorption spectra are shown for the formation of NADPH (λ_max_ = 340 nm) during a DEN2-NS3(S171-K618) reaction kinetics experiment at various time points: (i) 322 s, (ii) 282 s, (iii) 242 s, (iv) 202 s, (v) 162 s, (vi) 122 s, (vii) 82 s, and (viii) 41 s. The concentrations of the substrate and effector were as follows: substrate: [ATP] = 1.78 mM and effector [5’-ended overhang 31/18-mer dsDNA] = 17.9 μM. (b) A plot of absorbance at 340 nm (representing [NADPH]) vs. time (red trace) is presented for the same kinetics experiment as in (a). The tangent line that was made from the first five time points (R^2^ = 0.9995) showed a deviation from the plot and represented the initial velocity measurement of the reaction. The time optimum was determined to be 200 s. Data analysis and plots were prepared in GraphPad Prism.

The compound SSYA10-001, along with three catechin compounds EGCG, ECG, and EGC, were analyzed in a primary screen for inhibition against DEN2-NS3(S171-K618). K_i_ values and mode of inhibition were subsequently assessed. Briefly, reaction mixtures were carried out in a buffered solution [50 mM TEA (pH 7.6), 150 mM NaCl] and contained the following reagents at F.A.C.s: DEN2-NS3(S171-K618) (5.6 μg/mL), *Tl*GlcK (5.6 μg/mL), *Lm*G6PDH (5.6 μg/mL), NADP^+^ (0.477 mM), 5’-ended overhang 31/18-mer dsDNA (a putative effector at this concentration, 72.0 μM), MgCl_2_ (6.95 mM), DMSO (3.85% by vol.), D-glucose (1.75 mM), ATP (variable concentrations, *vide infra*), and inhibitor compound (variable concentrations, *vide infra*). The primary screening assay was performed with the substrate ATP at various concentrations of 40.0 μM, 80.0 μM, and 120.0 μM to produce Dixon and Cornish-Bowden plots having different slopes. Each of the inhibitor compounds was originally prepared at working concentrations of 10.0 mM and 1.0 mM, and these stocks were dissolved in 100% DMSO. The inhibitor F.A.C.s for EGCG and ECG ranged from 0.020 – 2.500 μM; the F.A.C. of SSYA10-001 ranged from 0.625 – 40.0 μM; and the F.A.C. for EGC ranged from 1.25 – 80.0 μM. The inhibitors were added to the reaction mixture (before initiation) for a minimum of 10 min. Each DEN2-NS3(S171-K618) reaction was performed at room temperature (25 °C) in a total volume of 200 µL. A 90-µL aliquot was then transferred to a black 96-well fluorescence microplate; this volume adjustment kept the NADPH fluorescence signal within the instrument’s linear detection range. The reaction was initiated by adding ATP and monitored continuously for an optimized duration of 200 s without termination. Without the use of a reaction-stopping solution, a timing adjustment was implemented for the microplate reader. For example, to run samples in row A ([Inh] = 40.0 µM), the microplate reader started at 2-min:48-s; for row B ([Inh] = 80.0 µM), the microplate reader started at 2-min:43-s; and for row C ([Inh] = 120.0 µM), the microplate reader started at 2-min:38-s. DEN2-NS3(S171-K618) activity in the presence of inhibitor was analyzed for the production of NADPH. The completed reaction mixtures were added to a black, round-bottom microplate and scanned by a VANTAstar-F multi-mode microplate reader. The instrument was set to fluorescence mode, and the wavelengths (λ_ex_ = 340 nm; λ_em_ = 485 nm) were set to observe the relative fluorescence units of NADPH. The determination of the K_i_ value and mode of inhibition for each inhibitor was assessed by using the Dixon and Cornish–Bowden plot analysis method [22]. Assays were performed in triplicate. A standard curve was implemented for NADPH (RFU vs. [NADPH]), in which the standard solutions of NADPH had a concentration range from 1.094 to 140.0 µM.

### 2.6. Confirmatory Assay

.The inhibitors SSYA10-001, EGCG, ECG, and EGC revealed an inhibitory effect in the presence of DEN2-NS3(S171-K618), *Tl*GlcK, and *Lm*G6PDH; however, further assessment was required to determine whether these compounds were indeed on-target confirmed hits of NTPase activity from DEN2-NS3(S171-K618). This was accomplished by removing DEN2-NS3(S171-K618) from the reaction and initiating the first enzyme of the enzyme – coupled system (*Tl*GlcK) using D-glucose and ADP as substrates. Compounds were typically tested as four trials (A1-A4) in a black, round-bottom, 96-well microplate (for fluorescence) in which wells were filled with a buffered solution [50 mM TEA (pH 7.6), 150 mM NaCl] amounting to a total volume of 200.0 μL, and contained the following reagents at final assay concentrations indicated: *Tl*GlcK (5.5 μg/mL), *Lm*G6PDH (5.5 μg/mL), NADP^+^ (0.477 mM), ADP (2.00 mM), MgCl_2_ (6.94 mM), DMSO (1.55% by vol.), D-glucose (1.75 mM), and inhibitor (the concentrations were 1.00 µM for EGCG, 10.0 µM for EGC, 0.80 µM for ECG, and 20.0 µM for SSYA10-001, which were selected based on being at least 4x greater than the determined K_i_ value for DEN2-NS3(S171-K618)). The inhibitor was pre-incubated with the *Tl*GlcK–*Lm*G6PDH enzyme-coupled system for at least 10 min prior to the assay. Reactions were initiated as described for the DEN2-NS3(S171-K618) assay. Before reading the plate with the microplate reader, a 90-µL aliquot from each 200 µL reaction mixture was transferred into a single well of a black 96-well fluorescence plate. NADPH was measured for its production using a VANTAstar microplate reader in fluorescence mode (λ_ex_ = 340 nm and λ_em_ = 485 nm) over 200 s. Each plate included eight positive controls (B1–B8) and eight negative controls (C1–C8). Positive controls matched the test samples, except we replaced the inhibitor with 100% DMSO. In the negative control samples, they were prepared by having the same reagents as the positive control samples, except that ADP and D-glucose were omitted (their volumes were replaced by buffer). The positive and negative controls would run in a 200-s incubation period at room temperature as well; however, since there was no reaction-stopping solution used in this experiment, a timing adjustment was implemented. For example, to run samples in row A (test reactions), the microplate reader was started at 2-min:48-s, for row B (positive controls), the microplate reader was started at 2-min:43-s, and for row C (negative controls), the microplate reader was started at 2-min:38-s. These samples were required for the calculation of normalized percent inhibition (NPI), where 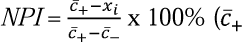 is the mean of the positive control, *C*_ is the mean of the negative control, and *x_i_* is the raw measurement on the i^th^ compound) [23]. Compounds were evaluated on whether they would inhibit the *Tl*GlcK – *Lm*G6PDH enzyme-coupled system by more than 25%. These results support that the observed inhibition was not due to interference with the *Tl*GlcK–*Lm*G6PDH coupled system and is consistent with DEN2-NS3(S171–K618)-directed inhibition.

### 2.7. In Silico Analysis: Protein Preparation, SiteMap-Guided Molecular Docking, MM-GBSA Rescoring, and Molecular Dynamics Simulations of DEN2-NS3(S171–K618)

The computational workflow was performed using Maestro 14.6 from the Schrödinger Suite (Schrödinger, LLC, New York, NY, USA). The crystallographic structure deposited in the Protein Data Bank under entry PDB_00002BHR [12] was used as the structural template for all *in silico* analyses. This structure corresponds to the DEN2-NS3 helicase catalytic region and is consistent with the recombinant DEN2-NS3(S171–K618) construct evaluated in the biochemical assays. Chain A was selected for protein preparation, binding-site prediction, molecular docking, MM-GBSA rescoring, and molecular dynamics simulations.

The protein structure was prepared using the Protein Preparation Wizard [24] implemented in Maestro. Bond orders were assigned, hydrogen atoms were added, and missing side-chain atoms were corrected when necessary. Protonation states were assigned using Epik [25] at pH 7.0 ± 2.0, and the hydrogen-bonding network was optimized. The prepared structure was then subjected to restrained energy minimization using the OPLS4 force field [26] to remove unfavorable contacts while preserving the crystallographic backbone geometry.

Potential ligand-binding regions were identified using SiteMap [27]. The docking site was not predefined as the canonical ATP-binding pocket. Instead, SiteMap was used to identify potentially druggable cavities in the prepared DEN2-NS3(S171-K618) helicase structure. The top-ranked pocket was selected for subsequent grid generation and docking because it showed favorable druggability and pocket-quality parameters, including a SiteScore, Dscore, size, and volume. The structural location of this SiteMap-selected pocket within the DEN2-NS3(S171-K618) helicase/NTPase domain is shown in **Figure S1** (see Supplementary Information). A receptor grid was generated around this selected pocket using a 20 Å grid box centered at X = 9.09, Y = 33.73, and Z = 48.88.

The ligand set included five catechin-related compounds and three reference molecules. Ligand structures were retrieved from PubChem using the following compound identifiers: (+)-catechin, CID 9064; (−)-epicatechin, CID 72276; (−)-epigallocatechin, CID 72277; (−)-epicatechin gallate, CID 107905; (−)-epigallocatechin gallate, CID 65064; ATP was included as a substrate/reference molecule [28], SSYA10-001 as a reference coronavirus nsp13 helicase inhibitor [29], and mosnodenvir as a DENV NS3-NS4B interaction inhibitor [30]. These reference molecules were used to contextualize the catechin docking results within the same predicted binding site. The ligands were prepared using LigPrep in Maestro. Ionization and tautomeric states were generated at pH 7.0 ± 2.0 using Epik, and ligand geometries were energy-minimized using the OPLS4 force field.

Molecular docking was performed using Glide in extra precision mode [31, 32]. All ligands were docked into the pocket using the same receptor grid. To assess numerical reproducibility of the docking/rescoring workflow, three repeated Glide XP docking/Prime MM-GBSA calculations were performed for each ligand using the same prepared receptor structure, receptor grid, ligand-preparation protocol, and scoring parameters. These runs were intended to assess procedural reproducibility and were not treated as independent conformational replicas. The resulting docking and MM-GBSA values were summarized as mean ± SD across the repeated calculations. The resulting SD values therefore reflect computational repeatability under the selected protocol rather than conformational or thermodynamic uncertainty. Reference molecules included ATP (substrate), SSYA10-001 (coronavirus nsp13 helicase inhibitor), and mosnodenvir (DENV NS3-NS4B interaction inhibitor). These reference molecules were used to contextualize the catechin docking results within the same predicted binding site. The corresponding 2D ligand–protein interaction diagrams for the catechins and reference compounds are provided in **Figure S2** (see Supplementary Information).

Post-docking energetic rescoring was performed using Prime MM-GBSA [33]. For each ligand, the top-ranked pose from each replicate docking run was rescored, and the resulting values were summarized as mean ± SD. Binding free-energy estimates were calculated using the equation ΔG_bind_ = G_complex_ − (G_protein_ + G_ligand_). These MM-GBSA values were used as relative end-point energetic descriptors for comparing ligand accommodation within the selected pocket and were not interpreted as direct experimental binding free energies.

Molecular dynamics simulations were carried out using Desmond to assess the dynamic stability of selected protein-ligand complexes [24]. Simulated systems included EGCG, ECG, ATP, mosnodenvir, and SSYA10-001 bound to the pocket of DEN2-NS3(S171-K618) helicase. Each complex was embedded in an orthorhombic simulation box filled with explicit TIP3P water molecules [34]. Counterions were added to neutralize the systems, and 0.15 M NaCl was added to reproduce physiological ionic strength. All systems were parameterized using the OPLS4 force field.

Before production simulations, each system was subjected to the default Desmond relaxation protocol, including restrained minimization and equilibration steps. Production molecular dynamics simulations were then performed under the NPT ensemble at 309.6 K for 200 ns. For each selected protein–ligand complex, three independent replicas were performed. The simulation trajectories were analyzed using the Simulation Interactions Diagram tool in Maestro. Protein stability was assessed by monitoring protein Cα root-mean-square deviation, protein root-mean-square fluctuation, and secondary structure stability. Ligand stability was evaluated using ligand RMSD relative to the protein, internal ligand RMSD, ligand RMSF, radius of gyration, molecular surface area, solvent-accessible surface area, polar surface area, and intramolecular hydrogen-bond formation. The three replicas were independently initialized and analyzed separately before replica-level summary statistics were calculated.

Protein–ligand interaction occupancies were calculated independently for each of the three 200-ns replicas. For each residue and interaction class, the fraction of trajectory frames containing the interaction was first calculated separately within each replica. Replica-level occupancies were then summarized as mean ± SD across the three independent trajectories. Individual replicate RMSD trajectories and replicate-level interaction occupancies are provided in the Supplementary Information.

## 3. Results and Discussion

### 3.1. Enzyme Kinetics Time Optimum Determination of DEN2-NS3(S171-K618)

The initial velocity measurement at the highest substrate concentration (300 nmol of ATP) for the assay was evaluated on its time optimum. Although fluorescence was used to monitor NADPH production in the enzymatic assays, UV–visible spectroscopy was implemented in order to measure NADPH production in the DEN2-NS3(S171-K618) NTPase activity assay for the time optimum. At room temperature and at F.A.C.s of 6.0 µg/mL each of DEN2-NS3(S171-K618), *Lm*G6PDH, and *Tl*GlcK, we conducted separate UV-visible scans at the following time points: 41, 82, 122, 162, 202, 242, 282, and 322 s (**Figure 3a**). The analysis of λ_max_ for NADPH (340 nm) as a function of time (**Figure 3b**) reveals a tangent line running through data points ranging from 41 to 202 s with an R^2^ value of 0.9995, and recorded time-points shown after 202 s indicated the loss of linearity. Thus, we determined the time optimum was 200 s.

The 5’-ended overhang 31/18-mer dsDNA originally appeared to slightly improve the NTPase activity of DEN2-NS3(S171-K618) relative to the basal reaction containing enzyme without nucleic acid. From a re-examination based on three duplex concentrations (5.0 µM, 20.0 µM, and 72.0 µM, **Table 1**), the effect was determined to be inconsistent in both magnitude and direction. When the duplex had a concentration of 5.0 µM, a mean NPI of 10.72 ± 4.59% was observed. This is a borderline loss in NTPase activity, which approached but did not reach significance (p = 0.056). Next, when the duplex had a concentration of 20.0 µM (a close representation of the concentration used in the time optimum experiment), a mean NPI of 5.25 ± 4.56% was observed, but was not distinct from the control (p = 0.184). At a duplex concentration of 72.0 µM (a representative concentration for the enzyme−inhibitor assays), the sign reversed where a mean NPI of −39.57 ± 20.65% was observed. This corresponded to 139.57% of NTPase activity and a trend toward allosteric activation (p = 0.080). Moreover, the replicate variance was large with respect to the mean, so with n = 3 these data do not carry much weighting to establish or dismiss an effect in either direction. Based on this, we report the duplex as a putative effector of undetermined direction rather than as an activator or an inhibitor. Finally, the 20-mer ssDNA at an F.A.C. of 5.0 µM performed as an inhibitor under the same conditions (NPI = 24.40 ± 0.86%, p = 0.0004). If residual unannealed ssDNA had been found in the 31/18-mer duplex preparation, we would expect a measurement bias to occur towards inhibition; however, at a concentration of 72.0 µM since we observed an approach towards activation, this rules out systematic ssDNA artifact contamination in the duplex preparation.

**Table 1.** Influence of ssDNA and dsDNA compounds on the activation and inhibition of DEN2-NS3(S171–K618) NTPase activity. ^a-c^.

| Condition | NPI (%),<br>mean ± SD | NTPase Activity (%),<br>mean ± SD |
| --- | --- | --- |
| Without ssDNA/dsDNA (Control) | 0.00 ± 0.00 | 100.00 ± 0.00 |
| 5.0 μM 20-mer ssDNA <sup>d</sup> | 24.40 ± 0.86 | 75.60 ± 0.86 <sup>e</sup> |
| 5.0 μM 31/18-mer 5'-overhang dsDNA | 10.72 ± 4.59 | 89.28 ± 4.59 <sup>f</sup> |
| 20.0 μM 31/18-mer 5'-overhang dsDNA | 5.25 ± 4.56 | 94.75 ± 4.56 <sup>g</sup> |
| 72.0 μM 31/18-mer 5'-overhang dsDNA | -39.57 ± 20.65 | 139.57 ± 20.65 <sup>h</sup> |

<sup>a</sup> Values are normalized percent inhibition (NPI) with respect to a reaction that does neither ssDNA nor dsDNA, which by definition has NPI = 0 and no associated variance. $NPI = \frac{\bar{c}_+ - x_i}{\bar{c}_+ - \bar{c}_-} \times 100\%$ , where $\bar{c}_+$ is the mean of the positive control, $\bar{c}_-$ is the mean of the negative control, and $x_i$ is the raw measurement on the $i^{th}$ condition [23]; and NTPase activity = 100% – NPI.
<sup>b</sup> Data are mean ± SD of three independent experiments ( $n = 3$ ).
<sup>c</sup> The p-values were calculated by one-sample t-test against the control (NPI = 0).
<sup>d</sup> The 20-mer ssDNA corresponds to 5'-AGAGTTTGATCCTGGCTCAG-3'.
<sup>e</sup> Weak inhibition is observed ( $p = 0.0004$ ).
<sup>f</sup> A borderline loss in NTPase activity is observed, approaching but not reaching statistical significance ( $p = 0.056$ ).
<sup>g</sup> No statistically significant inhibition is observed at this concentration ( $p = 0.184$ ).
<sup>h</sup> A trend toward allosteric activation was observed, but large replicate variance (SD of 20.65%) prevented a definitive statistical conclusion at this concentration ( $p = 0.080$ ).

### 3.2. Assessment of DEN2-NS3(S171-K618) Purity, Enzymatic Activity, and Inhibition

The protein purity of DEN2-NS3(S171-K618) and the enzyme-coupled system were evaluated by SDS-PAGE (**Figure 4a**). In particular, the analysis of His_6_-DEN2-NS3(S171-K618) helicase under reducing conditions, followed by staining with Coomassie Brilliant Blue R-250, revealed a major band corresponding to the monomeric subunit molecular mass of 52.8 kDa (shown in lane 2 of the figure). In comparison to the molecular weight standards, where there is a 50 kDa standard in lane 1, any slight migration discrepancy is ascribed to the inherent limitations of SDS-PAGE with regard to electrophoretic mobility. Lanes 3 and 4 of the same gel revealed the enzymes that are part of the enzyme-coupled system, which had single major bands corresponding to the monomeric subunit molecular masses of *Tl*GlcK (53.6 kDa, lane 3) and *Lm*G6PDH (54.4 kDa, lane 4).

**Figure 4:**
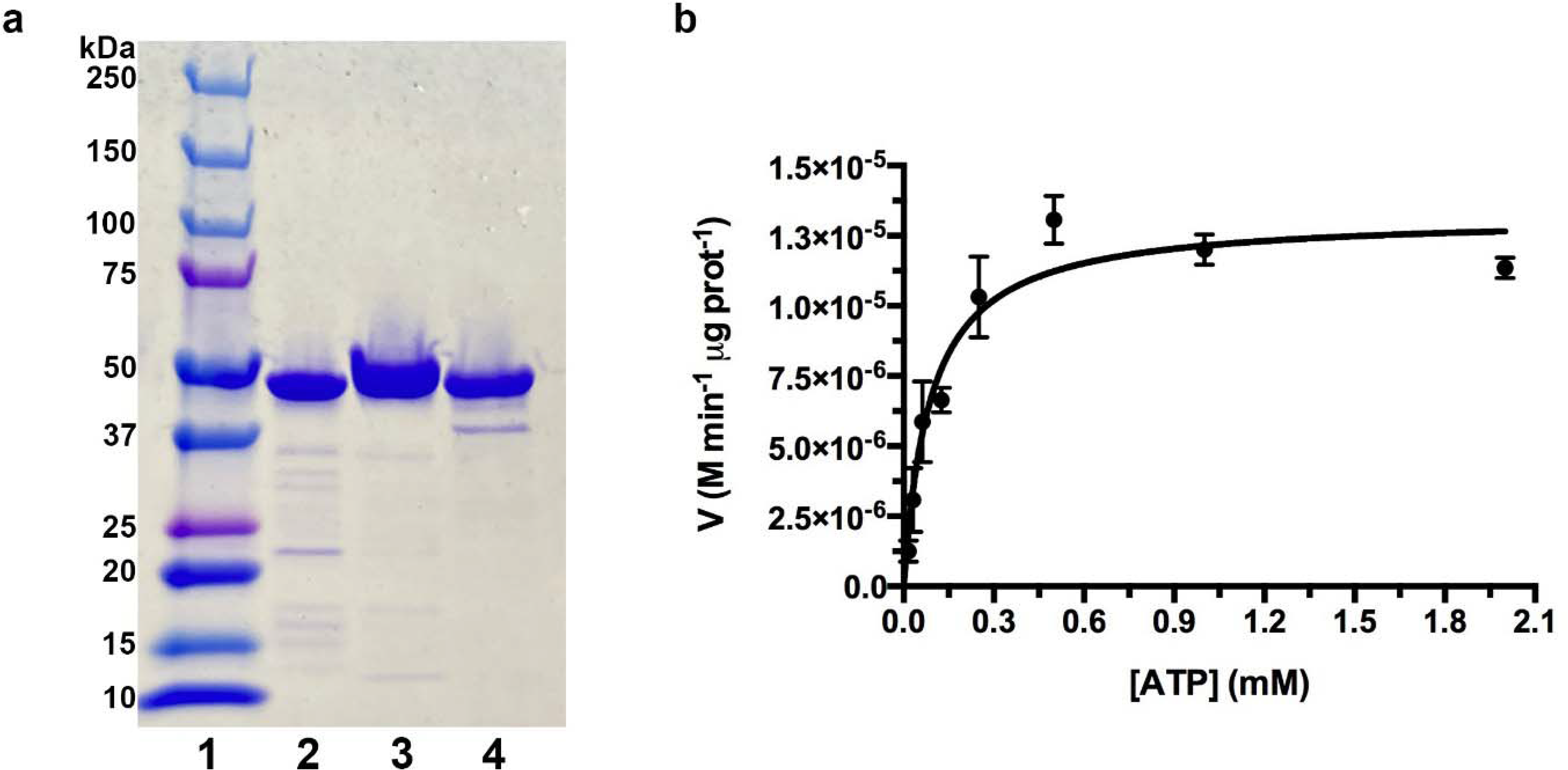
Enzyme purity assessment and fluorometric assay of DEN2-NS3(S171-K618). (a) SDS-PAGE of sample 1 representing the molecular weight marker ranging from 10 – 250 kDa; sample 2 representing DEN2-NS3(S171-K618) (the DENV-NS3 helicase/NTPase catalytic domain); GenBank accession no. WVS18090.1. Sample 3 represents *Thermococcus litoralis* (DSM 5473) ADP-dependent glucokinase (*Tl*GlcK); GenBank accession no. Q7M537.1. Sample 4 represents *Leuconostoc mesenteroides* glucose-6-phosphate dehydrogenase (*Lm*G6PDH); GenBank accession no. AAA25265.1. The monomeric subunit molecular masses for DEN2-NS3(S171-K618), *Tl*GlcK, and *Lm*G6PDH are 52,845 Da, 53,621 Da, and 54,441 Da, respectively. The 50 kDa standard (within sample 1) exhibited slightly different electrophoretic mobility compared to samples 2, 3, and 4. (b) Enzyme kinetics of DEN2-NS3(S171-K618) as represented by a Michaelis-Menten plot for the NTPase reaction. The plot shows the rate of NADPH formation (200-s time course) as a function of [ATP] (range of 0.0156 – 2.00 mM), in the presence of DEN2-NS3(S171-K618) and 72.0 µM 5’-overhang 31/18-mer dsDNA (where either the duplex or unannealed ssDNA acts as the effector). The K_M_ value for ATP was determined to be 88.6 ± 14.5 µM.

We report on the Michaelis-Menten parameters for DEN2-NS3(S171-K618) with respect to ATP as the substrate, for NTPase activity. We find a K_M_ of 88.6 ± 14.5 µM, a V_MAX_ of 1.322 x 10^-5^ ± 5.295 x 10^-7^ M min^-1^ μg prot^-1^, a k_cat_ of 125 ± 2 min^−1^, and a k_cat_/K_M_ of 1.41 ± 0.02 µM^−1^ min^−1^ (**Figure 4b**). The K_M_ value for DEN2-NS3(S171-K618) with respect to ATP was determined in this study to be 2.6-fold higher than a previously reported K_M_ value of 34 µM [12]. Four candidate inhibitors were then evaluated, such as SSYA10-001, EGCG, ECG, and EGC against DEN2-NS3(S171-K618) in a primary screen. The K_i_ values were determined by utilizing Dixon and Cornish-Bowden plots and one of three independent experiments are shown in **Figure 5**, whereas the reported K_i_ values correspond to the mean ± SD calculated from the three experiments, see **Table 2** and **Table S1** (Supplementary Information). Afterwards, confirmatory assays were performed from quadruplicate independent measurements, and the compounds were not observed in producing any significant *Lm*G6PDH inhibitory effects at the concentrations tested. The observed inhibitory constants and suggestive modes of inhibition (from Dixon and Cornish-Bowden plot analyses) of the tested compounds were as follows: K_i_ = 10.2 ± 0.3 μM for SSYA10-001 (mixed-mode inhibition), K_i_ = 400 ± 86.6 nM for EGCG (mixed-mode inhibition), K_i_ = 550 ± 250 nM for ECG (uncompetitive inhibition), and K_i_ = 18.3 ± 4.2 μM for EGC (mixed-mode inhibition, **Table 2**). Dixon and Cornish-Bowden plots are shown in **Figure 5** for each inhibitor compound, and the quantitative K_i_ values from individual trials can be found in **Table S1** (see Supplementary Information).

**Figure 5:**
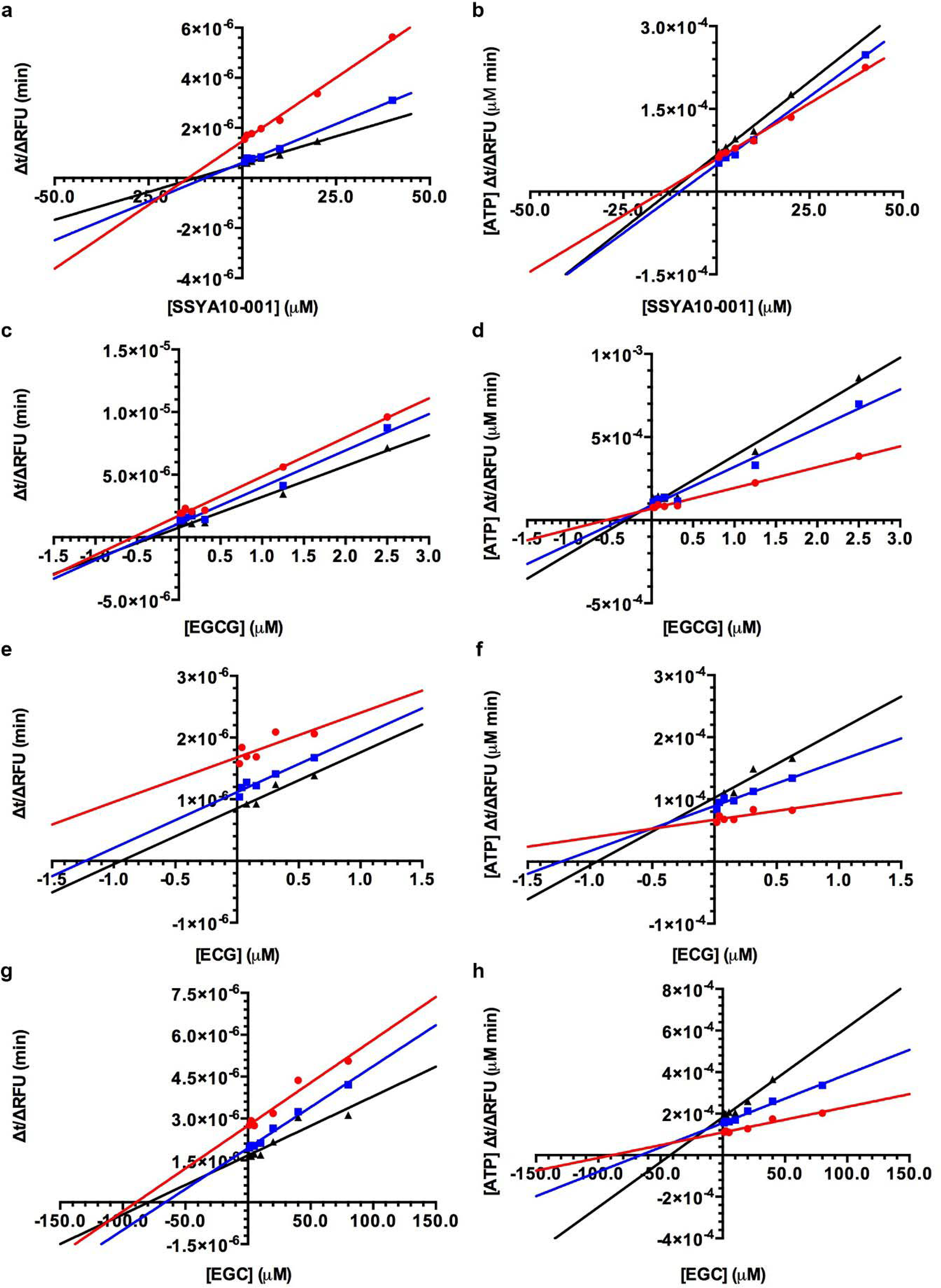
Enzyme-inhibitor kinetics for DEN2-NS3(S171-K618) and compounds SSYA10-001, EGCG, ECG, and EGC in order to determine K_i_ values and mode of inhibition. Panels (a, c, e, g) represent Dixon plots of inverse rate (Δt/ΔRFU) as a function of inhibitor concentration. Values calculated from the extrapolation of the intersection of the three lines with the X-axis were as follows: K_i_ = 10.2 ± 0.3 μM for SSYA10-001, K_i_ = 400 ± 86.6 nM for EGCG, K_i_ = 550 ± 250 nM for ECG, and K_i_ = 18.3 ± 4.2 μM for EGC. Panels (b, d, f, h) represent Cornish-Bowden plots of inverse rate multiplied by the substrate concentration ([ATP]*Δt/ΔRFU) as a function of inhibitor concentration. The modes of inhibition were determined as mixed-mode inhibition for SSYA10-001 and EGC; and potentially uncompetitive or mixed-mode inhibition for EGCG and ECG. In the other trials pertaining to EGCG and ECG, various Dixon plot lines would be observed (either having intersecting or parallel lines) and the assessment for these inhibitor binding modes should be further evaluated, experimentally. The inhibitor concentrations ranged from 0.0–40.0 μM for SSYA10-001, 0.0–2.5 μM for EGCG, 0.0–0.625 μM for ECG, and 0.0–80.0 μM for EGC. For all inhibitors examined, three different substrate concentrations were analyzed, 40 μM ATP (red circles), 80 μM ATP (blue squares), and 120 μM ATP (black triangles). Dixon and Cornish–Bowden plots from one of three independent experiments are shown. Quantitative reproducibility is reflected in the K_i_ values reported as mean ± SD, see **Table 2** and **Table S1** (Supplementary Information). Dixon plot and Cornish-Bowden plot graphs corresponding to an individual compound are from the same experiment.

**Table 2.**
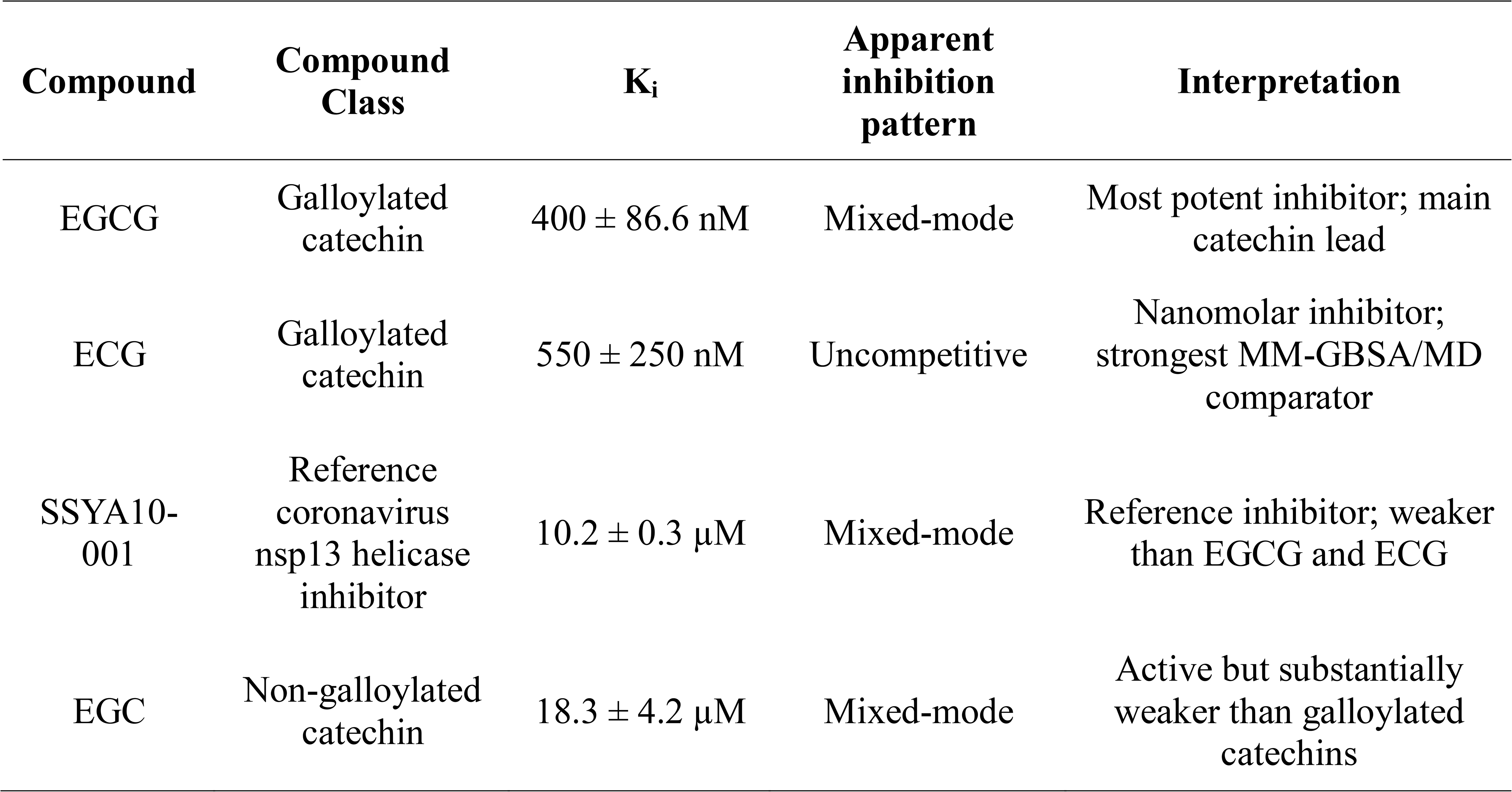
Inhibition constants and proposed modes of inhibition for catechin compounds and SSYA10-001 against DEN2-NS3(S171–K618).

| Compound | Compound Class | K<sub>i</sub> | Apparent inhibition pattern | Interpretation |
| --- | --- | --- | --- | --- |
| EGCG | Galloylated catechin | 400 ± 86.6 nM | Mixed-mode | Most potent inhibitor; main catechin lead |
| ECG | Galloylated catechin | 550 ± 250 nM | Uncompetitive | Nanomolar inhibitor; strongest MM-GBSA/MD comparator |
| SSYA10-001 | Reference coronavirus nsp13 helicase inhibitor | 10.2 ± 0.3 µM | Mixed-mode | Reference inhibitor; weaker than EGCG and ECG |
| EGC | Non-galloylated catechin | 18.3 ± 4.2 µM | Mixed-mode | Active but substantially weaker than galloylated catechins |

The relative standard deviation for the K_i_ value with respect to the inhibitor SSYA10-001 was 2.94%. This was in sharp contrast to the tightly binding catechin inhibitors EGCG and ECG that exhibited relative standard deviations in K_i_ of 21.7% and 45.5%, respectively. EGC also revealed a high relative standard deviation in K_i_ of 23.0%. Since the reference inhibitor had a very low relative standard deviation, we can rule out systemic deviations such as microplate reader timing variations during data acquisition and/or cumulative pipetting errors, as there indeed were numerous pipetting steps required for the reaction setup. Instead, we attribute the property of photolability as the likely problematic issue to our assay with respect to the three test inhibitors, which all belong to the catechin scaffold. It is likely that light from inside the laboratory may have decomposed the catechin inhibitors during assay preparation, which resulted in uneven concentration shifts, ultimately leading to the spike in the standard deviation. Catechin compounds have been documented to be light sensitive, and the photodegradation of EGCG was studied by Bianchi and colleagues. In their irradiation studies, EGCG was found to undergo a loss of 68.9% after being exposed to 1 h of natural sunlight [35]. Thus, it is plausible that some photodegradation took place during the setup of the assay.

The distinction between mixed-mode and uncompetitive inhibition in the trials pertaining to the galloylated catechin inhibitors EGCG and ECG, respectively (through analysis of their Dixon and Cornish-Bowden plots) is difficult to firmly establish. This ambiguity arises since only the Dixon plots reveal parallel or intersecting lines between trials. We recommend an orthogonal experimental, biophysical technique to explore the inhibition mode of these catechin compounds for future studies. In particular, isothermal titration calorimetry can directly monitor physical binding of the galloylated catechins to the free enzyme vs. the enzyme-substrate complex. For the determination of uncompetitive inhibition, the titration of the catechin inhibitor directly into free enzyme (DEN2-NS3) will yield no heat release because uncompetitive inhibitors cannot bind to the free enzyme. However, if the titration of the catechin compound is performed with DEN2-NS3 that is pre-incubated with a non-hydrolyzable compound (in replacement of the substrate ATP), such as AMP-PNP, then the heat of binding will instead be generated. In the determination of mixed-mode inhibition by this technique, this would involve the titration of the galloylated catechin inhibitor into the free enzyme to produce a distinct heat of binding so that a binding constant can be established. If a second titration is performed (as a confirmatory test), where AMP-PNP is pre-incubated with DEN2-NS3, it is expected that a shifted binding affinity will be observed, and confirming that the inhibitor can indeed bind to both states.

### 3.3. Computational prioritization of EGCG and ECG within an amphipathic pocket

To complement the biochemical inhibition data, a structure-based computational workflow was performed to rationalize the interaction of catechin-derived compounds with the DEN2-NS3(S171–K618) helicase/NTPase catalytic domain. Experimentally, EGCG was identified as the most active inhibitor, with a K_i_ value of 400 ± 86.6 nM, followed closely by ECG with a K_i_ of 550 ± 250 nM. In contrast, the reference coronavirus nsp13 helicase inhibitor SSYA10-001 and the non-galloylated catechin EGC showed weaker inhibition, with K_i_ values of 10.2 ± 0.3 µM and 18.3 ± 4.2 µM, respectively (**Table 2**). These results indicate that the galloylated catechins EGCG and ECG are the strongest biochemical inhibitors of DEN2-NS3(S171–K618), while also suggesting that inhibitory potency is influenced by both catechin substitution pattern and binding mode.

The computational analysis was based on the crystallographic DEN2-NS3 helicase/NTPase catalytic domain (entry PDB_00002BHR), which corresponds to the S171–K618 region evaluated experimentally [12]. This domain is structurally relevant because flavivirus NS3 helicases couple NTP hydrolysis to nucleic acid binding and unwinding [36]. Previous structural work on DEN2-NS3 helicase/NTPase showed that this catalytic region contains conserved motifs involved in NTP binding and hydrolysis, as well as a positively charged tunnel compatible with single-stranded nucleic acid accommodation [12]. Thus, ligand binding within this domain may interfere with helicase function by perturbing NTPase coupling, stabilizing nonproductive conformations, or disrupting RNA-binding/translocation-associated motions.

In the present workflow, the docking site was not manually imposed as the crystallographic ATP-binding site. Instead, SiteMap was used to identify a druggable cavity in the prepared DEN2-NS3(S171-K618) helicase/NTPase structure. The selected pocket showed favorable druggability parameters, including a SiteScore of 1.120, Dscore of 1.042, and volume of 771.750 Å³. The pocket also displayed an amphipathic profile, featuring a hydrophilic score of 1.303 and a hydrophobic score of 0.618. The complete SiteMap descriptors, including size, enclosure, exposure, donor/acceptor character, and hydrophilic/hydrophobic scores, are provided in **Table S2** (see Supplementary Information). The spatial location and close-up view of this pocket are shown in **Figure S1** (see Supplementary Information). Overall, these descriptors support the selection of a large and polar pocket compatible with polyhydroxylated catechins capable of forming hydrogen bonds and water-mediated interactions.

The overall *in silico* workflow, representative EGCG/ECG binding poses, computational ranking, ligand RMSD profiles, and residue interaction persistence are summarized in **Figure 6**. The docking and MM-GBSA values are reported as mean ± SD from three repeated docking/rescoring calculations, whereas the binding poses shown in **Figure 6b** correspond to representative poses selected for structural interpretation. The docking results supported EGCG as the strongest catechin candidate at the initial screening level. EGCG showed the most favorable Glide XP docking score among all evaluated compounds, −9.914 ± 0.85, followed by ECG at −8.679 ± 0.42 and ATP at −8.088 ± 0.02 (**Table 3** and **Figure 6c**). The lower docking scores of EGC, catechin, and epicatechin suggest that galloylated catechins are better accommodated within the selected pocket. This trend is chemically reasonable because the gallate ester expands the interaction surface of the catechin scaffold and introduces additional hydroxyl groups capable of forming polar contacts. Thus, the docking results are consistent with the experimental observation that EGCG was the most potent inhibitor in the biochemical assay, while also supporting ECG as a closely related galloylated catechin lead.

**Figure 6:**
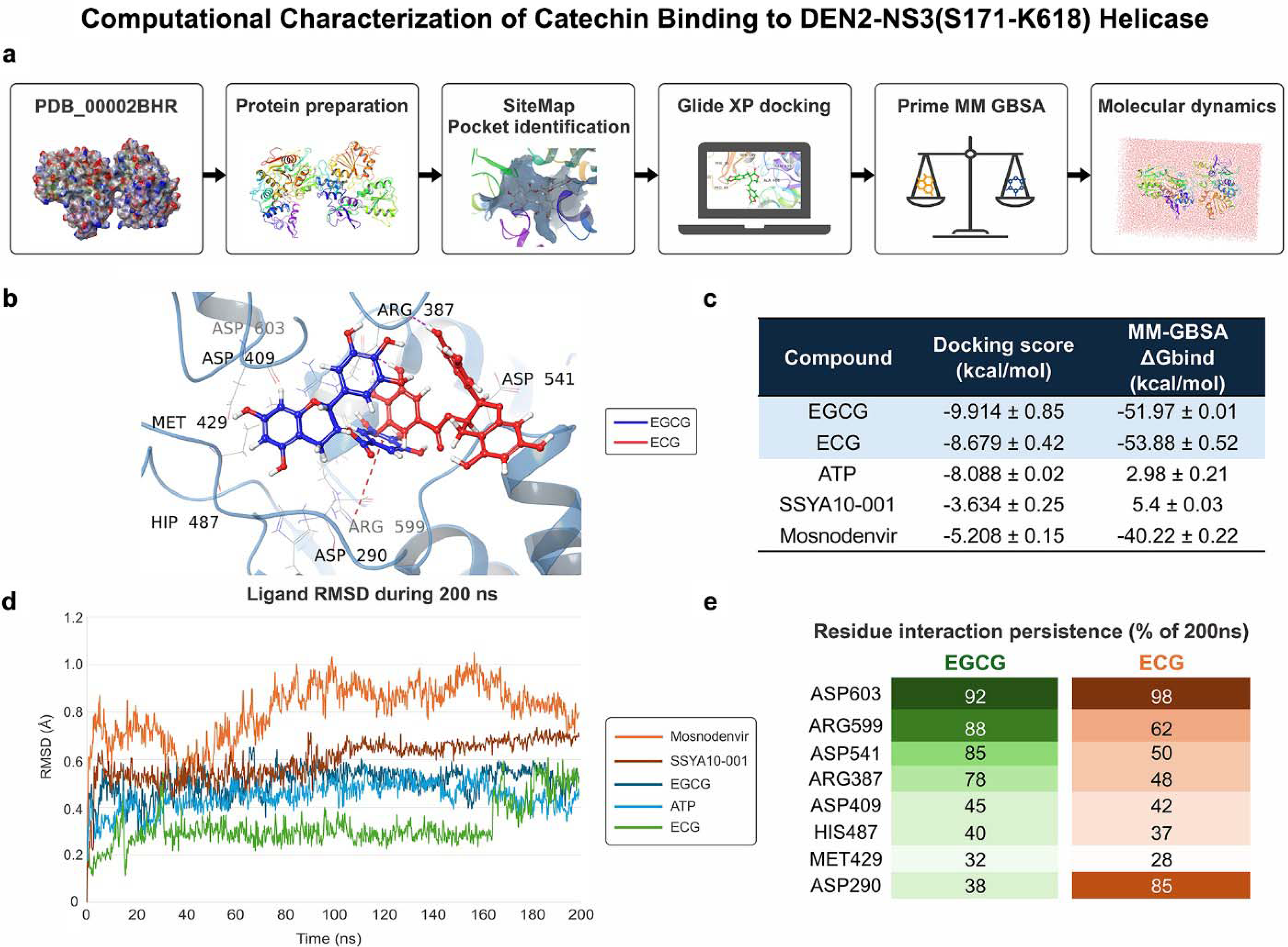
Computational characterization of catechin binding to DEN2-NS3(S171-K618) helicase. (a) *In silico* workflow for DEN2-NS3(S171–K618), including protein preparation from entry PDB_00002BHR, SiteMap pocket identification, Glide XP docking, Prime MM-GBSA rescoring, and 200 ns molecular dynamics simulations. (b) Representative EGCG and ECG binding poses within the amphipathic pocket. (c) Computational ranking of selected ligands by Glide XP docking score and Prime MM-GBSA ΔG_bind_ estimates; values are reported as mean ± SD across three independent replicate docking/rescoring calculations. (d) Representative ligand RMSD profiles relative to the protein-aligned frame during one of the three independent 200-ns simulations. Individual RMSD trajectories from all three replicas are provided in Figure S3. (e) Residue interaction persistence for EGCG and ECG; replica-level occupancies were calculated independently and summarized across the three trajectories as mean ± SD, with individual values provided in Table S5.

**Table 3.**
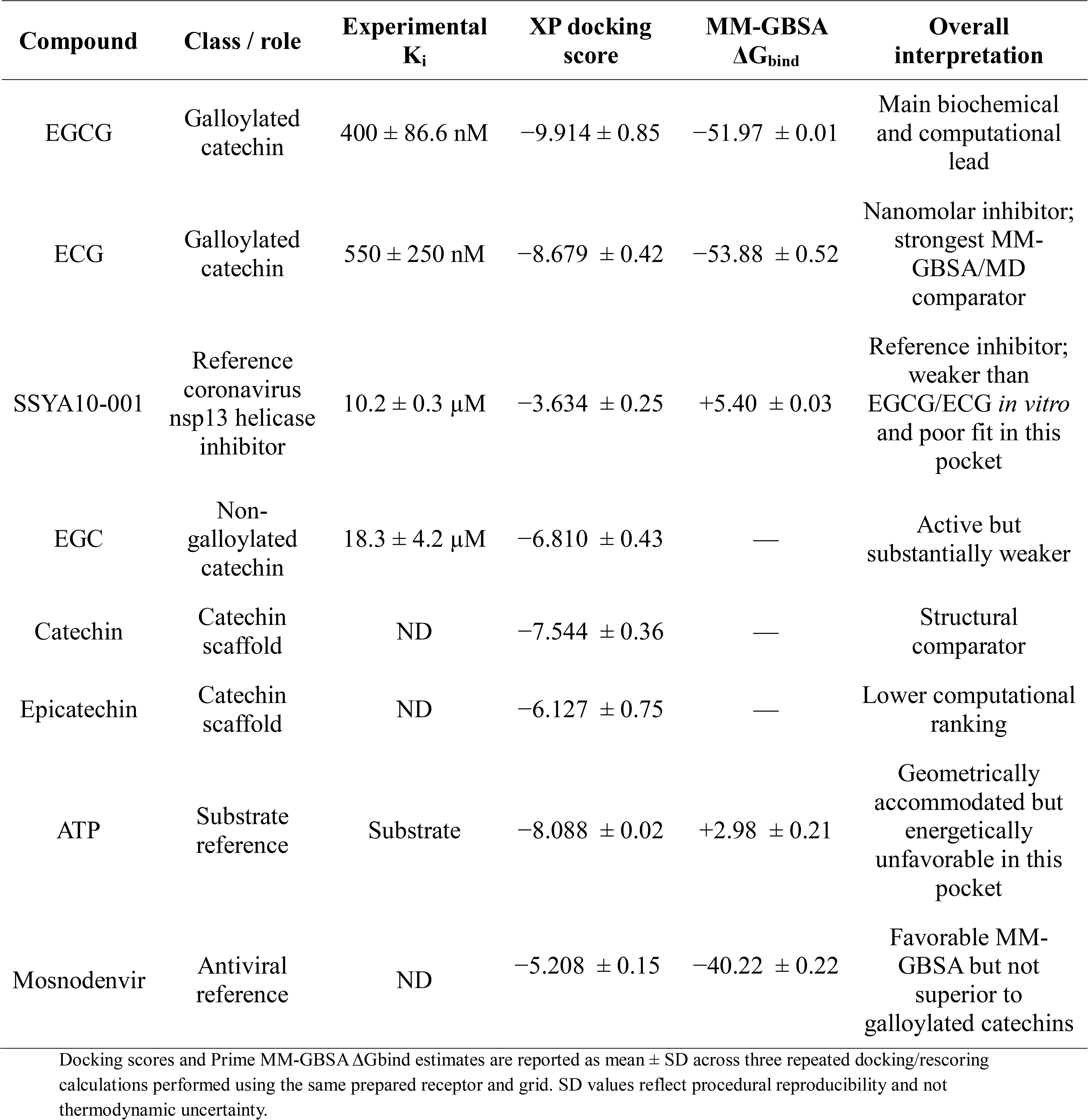
Experimental inhibition and computational ranking of DEN2-NS3(S171-K618) inhibitors.

| Compound | Class / role | Experimental $K_i$ | XP docking score | MM-GBSA $\Delta G_{\text{bind}}$ | Overall interpretation |
| --- | --- | --- | --- | --- | --- |
| EGCG | Galloylated catechin | $400 \pm 86.6$ nM | $-9.914 \pm 0.85$ | $-51.97 \pm 0.01$ | Main biochemical and computational lead |
| ECG | Galloylated catechin | $550 \pm 250$ nM | $-8.679 \pm 0.42$ | $-53.88 \pm 0.52$ | Nanomolar inhibitor; strongest MM-GBSA/MD comparator |
| SSYA10-001 | Reference coronavirus nsp13 helicase inhibitor | $10.2 \pm 0.3$ μM | $-3.634 \pm 0.25$ | $+5.40 \pm 0.03$ | Reference inhibitor; weaker than EGCG/ECG *in vitro* and poor fit in this pocket |
| EGC | Non-galloylated catechin | $18.3 \pm 4.2$ μM | $-6.810 \pm 0.43$ | — | Active but substantially weaker |
| Catechin | Catechin scaffold | ND | $-7.544 \pm 0.36$ | — | Structural comparator |
| Epicatechin | Catechin scaffold | ND | $-6.127 \pm 0.75$ | — | Lower computational ranking |
| ATP | Substrate reference | Substrate | $-8.088 \pm 0.02$ | $+2.98 \pm 0.21$ | Geometrically accommodated but energetically unfavorable in this pocket |
| Mosnodenvir | Antiviral reference | ND | $-5.208 \pm 0.15$ | $-40.22 \pm 0.22$ | Favorable MM-GBSA but not superior to galloylated catechins |

Docking scores and Prime MM-GBSA $\Delta G_{\text{bind}}$ estimates are reported as mean ± SD across three repeated docking/rescoring calculations performed using the same prepared receptor and grid. SD values reflect procedural reproducibility and not thermodynamic uncertainty.

Prime MM-GBSA rescoring provided a relative end-point energetic assessment of ligand accommodation within the selected pocket. ECG showed the most favorable ΔG_bind_ estimate, −53.88 ± 0.52 kcal/mol, followed closely by EGCG at −51.97 ± 0.01 kcal/mol (**Table 3** and **Figure 6c**). These values should be interpreted as comparative computational descriptors rather than direct estimates of experimental binding free energy. The complete energetic decomposition of the MM-GBSA calculations, including coulombic, covalent, hydrogen-bonding, lipophilic, solvation, and van der Waals components, is provided in **Table S3** (see Supplementary Information). The small difference between EGCG and ECG suggests that both galloylated catechins are energetically comparable within the selected pocket. Although ECG showed the most favorable MM-GBSA binding free energy and was also a nanomolar inhibitor, EGCG remained slightly more potent experimentally and showed the best XP docking score. This indicates that both ECG and EGCG are strong galloylated catechin leads, with EGCG providing the best overall biochemical–computational consensus and ECG providing the strongest energetic and MD-stability comparator.

The reference molecules provided a useful mechanistic context. ATP showed a relatively favorable XP docking score but an unfavorable MM-GBSA ΔG_bind_ of +2.98 ± 0.21 kcal/mol (**Table 3** and **Table S3** (see Supplementary Information)), suggesting that although ATP can geometrically occupy the selected pocket and form polar contacts, the energetic balance in this SiteMap-defined cavity is not favorable. Thus, ATP should be interpreted as a substrate/reference molecule rather than as evidence that the selected pocket corresponds to the canonical ATP-binding site. Although SSYA10-001 served as the reference coronavirus nsp13 helicase inhibitor, it was less potent than the galloylated catechins EGCG and ECG in the biochemical assay. SSYA10-001 showed mixed-mode inhibition with a K_i_ of 10.2 ± 0.3 µM, whereas EGCG and ECG displayed nanomolar K_i_ values. Computationally, SSYA10-001 also showed weak compatibility with the selected pocket, with a docking score of −3.634 ± 0.25 kcal/mol and an unfavorable MM-GBSA ΔG_bind_ of +5.40 ± 0.03 kcal/mol (**Table 3** and **Table S3** (see Supplementary Information)). This suggests that EGCG and SSYA10-001 may inhibit DEN2-NS3(S171-K618) helicase through different binding modes or distinct regions of the helicase/NTPase domain. Mosnodenvir, included as a DENV NS3-NS4B interaction inhibitor, showed a favorable MM-GBSA value but did not outperform EGCG or ECG, supporting its role as a contextual antiviral reference rather than a direct DEN2-NS3(S171-K618) helicase-site comparator. Representative 2D ligand–protein interaction diagrams for the catechin derivatives and reference compounds are provided in **Figure S2** (see Supplementary Information).

### 3.4. Molecular dynamics and mechanistic interpretation of catechin inhibition

Molecular dynamics simulations provided additional exploratory insight into the behavior of selected ligands within the SiteMap-selected pocket. Across the analyzed trajectories, protein Cα RMSD profiles reached system-dependent plateaus, while the ligand RMSD profiles revealed distinct degrees of local ligand displacement and reorientation.. EGCG and ECG showed comparatively lower ligand conformational drift than the reference compounds across the replicate analyses, supporting local ligand retention within the selected pocket.. This behavior is consistent with previous flaviviral NS3 helicase modeling studies in which 100 ns MD simulations were used to assess RMSD/RMSF profiles, radius of gyration, hydrogen-bond persistence, and ligand–protein interaction stability [11,37]. Therefore, the present MD results are interpreted as supportive and hypothesis-generating, rather than as exhaustive conformational sampling.

ECG displayed a stable local interaction profile across the MD analyses, maintaining recurrent contacts with ASP603 and ASP290, along with additional interactions involving ARG387, ASP541, ARG599, GLU490, VAL544, HIS487, and LYS388 (**Table 4** and **Table S4** (see Supplementary Information)). ASP603 was especially important, forming highly persistent hydrogen-bond interactions with ECG throughout the trajectory. This interaction pattern helps explain why ECG achieved the most favorable MM-GBSA energy and a stable MD profile. Although ECG showed strong binding stability and the most favorable MM-GBSA value, EGCG remained slightly more potent in the biochemical assay. This suggests that both compounds are highly relevant galloylated catechin leads, but that productive inhibition depends not only on energetic stability within the pocket, but also on how ligand binding perturbs the catalytic and conformational coupling of the helicase/NTPase domain.

**Table 4.** Molecular dynamics stability summary of selected DEN2-NS3(S171-K618) ligand complexes.

| Compound | Main MD behavior | Key residues | Interpretation |
| --- | --- | --- | --- |
| EGCG | Stable interaction network, moderate ligand mobility | ARG599, ASP541, ASP603, ARG387, MET429 | Best experimental-computational consensus |
| ECG | Strong dynamic stability and persistent H-bonds | ASP603, ASP290, ARG387, ASP541, ARG599, VAL544 | Best MM-GBSA/MD comparator |
| ATP | Many contacts but unfavorable MM-GBSA | ARG387, LYS388, THR408, ASP409, HIS487, ARG599, ASP603 | Not optimized for this SiteMap pocket |
| Mosnodenvir | Reorientation during MD | ASP409, CYS428, MET429, ILE365, HIS487 | Contextual antiviral reference |
| SSYA10-001 | Displacement/reorientation, limited favorable energetics | HIS487, ARG599, ASP409, ARG387, CYS485 | Reference inhibitor does not validate this pocket |

EGCG showed the strongest overall agreement between the biochemical inhibition data and computational ranking. It was the most potent inhibitor in the enzyme assay, displayed the best XP docking score, retained a favorable MM-GBSA energetic profile, and maintained a recurrent interaction involving ARG599, ASP541, ASP603, ARG387, MET429, SER602, and neighboring residues (**Table 4** and **Table S4** (see Supplementary Information)). This broader polar network may be more compatible with functional perturbation of the helicase/NTPase domain, particularly in light of the suggestive mixed-mode inhibition observed experimentally. EGCG therefore appears to provide the best balance between binding strength, pocket compatibility, and biochemical inhibition. This interpretation is consistent with previous work on ZIKV-NS3 helicase/NTPase, where EGCG was proposed to interact with ATPase-associated and RNA-binding regions and to inhibit NTPase activity [11].

The recurring residues observed in the MD simulations suggest that the selected pocket corresponds to a polar interaction hub within the DEN2-NS3(S171-K618) helicase/NTPase domain. ARG599, ASP603, ASP541, ARG387, LYS388, ASP409, HIS487, and MET429 appeared repeatedly throughout catechin and reference simulations (**Table 4** and **Table S4** (see Supplementary Information)). Among these, ASP603 and ARG599 were particularly relevant for catechin binding, whereas ASP409, HIS487, and MET429 appeared to define a shared pocket region also contacted by reference compounds. The shared interaction map provided in **Table S4** (see Supplementary Information) further highlights that EGCG and ECG converge on a partially overlapping residue network, while ATP, SSYA10-001, and mosnodenvir show distinct interaction patterns within the same region. Because helicase activity depends on coordinated motions between NTPase motifs, RNA-binding surfaces, and domain rearrangements [38], stabilization of this pocket by galloylated catechins could interfere with the conformational transitions required for efficient NTP hydrolysis and nucleic acid unwinding.

This interpretation is consistent with the inhibition pattern observed experimentally. The biochemical assay monitored DEN2-NS3(S171–K618) helicase/NTPase function through ATP-dependent generation of ADP and coupled NADPH formation, building on prior enzyme-coupled helicase assay development [19]. EGCG inhibited this activity more potently than SSYA10-001, with a nanomolar K_i_ value and mixed-mode inhibition (**Table 2**). However, the computational results do not require EGCG to occupy the canonical ATP-binding site. Instead, they support the possibility that EGCG binds a nearby or functionally connected amphipathic pocket capable of indirectly modulating enzyme function. This interpretation is compatible with mixed-mode inhibition, which may arise from ligand effects on both substrate-associated and conformational components of helicase activity.

Previous DENV cell-based studies showed that EGCG and ECG can inhibit DENV infection, with EGCG showing stronger dose-dependent antiviral activity [10]. Those studies proposed that EGCG can act early during infection, likely by interacting with virions and impairing viral entry. The nanomolar inhibition observed here for EGCG and ECG against DEN2-NS3(S171–K618), therefore, extends previous cell-based antiviral observations by identifying the helicase/NTPase catalytic domain as a plausible enzymatic target for galloylated catechins. The present work does not contradict an entry-related mechanism; rather, it expands on the possible antiviral profile of EGCG by showing that it can also inhibit the DEN2-NS3(S171-K618) helicase/NTPase catalytic domain *in vitro*.

The comparison with prior ZIKV-NS3 helicase/NTPase work is especially relevant. Kumar and colleagues reported that EGCG inhibited ZIKV-NS3 NTPase activity and used docking and MD simulations to propose interactions at both ATPase and RNA-binding regions [11]. Although residue numbering differs between ZIKV and DENV, the conceptual parallel is important: EGCG appears capable of engaging polar and dynamic regions of flaviviral helicases that are relevant to NTPase/RNA-binding coupling. The repeated involvement of charged and polar residues such as ARG599, ASP603, ARG387, ASP409, and HIS487 in the present DEN2-NS3(S171-K618) simulations supports a related mechanism in which EGCG binding stabilizes an amphipathic pocket and potentially alters helicase dynamics (**Table 4** and **Table S4** (see Supplementary Information)).

The results can also be interpreted in the context of previous work describing epigallocatechin derivatives as flaviviral protein-binding scaffolds. Coronado et al. showed that epigallocatechin-type molecules can inhibit flaviviral NS2B/NS3 proteases and may act through more than one inhibitory mode depending on the compound and viral target [18]. Although that work focused on the protease domain rather than the helicase/NTPase domain, it supports the broader concept that catechin-derived molecules can engage flaviviral nonstructural proteins through diverse interaction modes. The present results extend this concept to DEN2-NS3(S171-K618) helicase/NTPase inhibition.

Overall, the integrated data support EGCG as the principal lead for DEN2-NS3(S171-K618) helicase inhibition. EGCG was the most potent inhibitor in the enzyme assay, showed mixed-mode inhibition, displayed the best XP docking score, retained a strongly favorable MM-GBSA binding energy, and maintained persistent interactions with the amphipathic pocket during MD (**Tables 1–3** and **Tables S1–S3** (see Supplementary Information)). ECG should also be emphasized as a second nanomolar galloylated catechin lead because it showed strong biochemical inhibition, the most favorable MM-GBSA energy, and the most stable MD interaction profile. Thus, EGCG and ECG highlight the importance of galloylated catechin recognition in this pocket, whereas EGC, catechin, and epicatechin provide structure–activity context for weaker or lower-ranked catechin scaffolds. The galloylated catechins, particularly EGCG and ECG, inhibit DEN2-NS3(S171-K618) through interaction with a large amphipathic pocket within the helicase/NTPase domain. This pocket is distinct from the crystallographic ATP-binding site; rather, it is a druggable cavity composed of residues that form the functional RNA-binding channel. This is evidenced by the finding of persistent interactions with residues ARG599, ASP603, ASP541, ARG387, ASP409, HIS487, and MET429, which provide a structural rationale for catechin stabilization in this region [39]. These findings support EGCG as the main catechin-derived lead and ECG as a strong nanomolar comparator for future optimization of galloylated polyphenols as flaviviral helicase-targeting compounds.

The results should be interpreted as a mechanistic model rather than definitive proof of the exact binding site. Future studies should evaluate alternative pockets, compare the site against canonical NTPase and RNA-binding regions, and test site-directed mutants of residues such as ARG599, ASP603, ASP541, ARG387, ASP409, and HIS487. Biochemical assays evaluating ATP competition, nucleic-acid substrate dependence, and direct binding would further clarify whether EGCG acts through allosteric inhibition, RNA-binding interference, or partial overlap with NTPase-coupled motions.

## 4. Conclusions

Three catechins naturally derived from *Camellia sinensis* were tested against the DEN2-NS3(S171-K618) helicase/NTPase catalytic domain: the free catechin EGC and the galloylated catechins EGCG and ECG. In terms of the general structural scaffold, EGC differs from EGCG and ECG by the absence of a galloyl ester moiety at the C-3 position. This difference was enough to cause a major drop in inhibitor potency, where EGC revealed a K_i_ of 18.3 ± 4.2 µM. EGCG and ECG showed potencies that were 45.8-fold and 33.3-fold stronger than EGC, respectively (**Figure 5**). Consequently, we prioritized the galloylated catechins for subsequent in-depth computational studies. In our computational workflow and assessment, SiteMap [27] was utilized to find druggable cavities on DEN2-NS3(S171-K618) for the potent catechin inhibitors EGCG and ECG. One pocket identified was lined with residues ASP603, ARG599, ASP541, ARG387, ASP409, HIS487, MET429, and ASP290, which are residues that occupy the RNA-binding cavity (**Figure 6b** and **Figure S1** (Supplementary Information)) [12,40]. Subsequent exploratory MD simulations indicated that these catechin inhibitors showed stable local retention within the predicted pocket over the analyzed MD timescale.. This stability was evidenced by persistent amino acid sidechain contacts (**Figure 6e**). In particular, the sidechains from ASP603, ARG599, ASP541, and ARG387 formed shared, highly persistent contacts with over 45% occupancy throughout the 200-ns simulation trajectory. The protein Cα RMSD profiles showed system-dependent stabilization over the analyzed trajectories, whereas EGCG and ECG exhibited comparatively lower ligand conformational drift than the reference compounds, consistent with greater local retention within the selected pocket over the 200-ns timescale. Since these conserved residues normally dictate the electrostatic coordination of the viral ssRNA phosphate backbone within the helicase domain [12], the high occupancy observed during our MD simulations suggest that these catechins may interfere with the RNA-binding assembly [39]. Since EGCG/ECG binding occurs outside the catalytic NTPase region, an allosteric mode of inhibition is proposed rather than direct competition with ATP.

By complementing the experimental enzyme-inhibitor kinetics data (K_i_ values and suggestive modes of inhibition), the computational results support our proposed model interaction within this binding pocket. These data revealed a clear consistency with the target binding pocket location (identified by SiteMap) and the mixed-mode/uncompetitive modes of inhibition derived from our Dixon/Cornish-Bowden plot analyses with respect to the NTPase binding cavity (**Figure 5**). In agreement with the experimental potency rankings (K_i_ values of 400 nM for EGCG and 550 nM for ECG), molecular docking predicted highly favorable docking scores (−9.914 kcal/mol for EGCG and -8.679 kcal/mol for ECG). Accordingly, the highly favorable MM-GBSA profiles of EGCG and ECG provide relative energetic support for ligand accommodation within the selected pocket, but they should not be interpreted as evidence of a confirmed thermodynamic driving force. Future studies using microsecond-scale MD simulations, additional independent replicas, site-directed mutagenesis, and direct binding assays will be required to validate the proposed binding model and clarify the functional consequences of catechin binding.

While EGCG and ECG exhibit potent binding within the Dengue virus RNA-binding cavity, which was verified by the enzyme-coupled confirmatory assay (see **Figure 2b**, Eqs. 2b-3b), the progression of these specific compounds is hindered by well-documented liabilities. In particular, EGCG is classified as a pan-assay interference compound due to its target promiscuity [41], light-sensitive hydroxyl groups [35], and the tendency to induce hydrogen-bond-mediated protein aggregation [42]. To overcome these limitations, future medicinal chemistry optimization of the EGCG scaffold should focus on designing derivatives that mask or modify these problematic hydroxyl groups. This can be accomplished by their removal or by rendering them as prodrug moieties (e.g., via esterification) [43,44]. Such structural modifications are expected to improve metabolic and chemical stability while eliminating promiscuity-related protein-binding interference.

## CRediT authorship contribution statement

**Milla K. Wojciechowski:** Data collection, Formal analysis, Writing – Original draft, Review & Editing. **Luis Daniel Goyzueta-Mamani:** Conceptualization, Software, Data collection, Formal analysis, Writing – Original draft, Review & Editing. **Miguel Angel Chávez-Fumagalli:** Formal analysis, Writing – Review & Editing. **Edward L. D’Antonio:** Conceptualization, Project administration, Supervision, Funding acquisition, Formal Analysis, Validation, Writing – Original draft, Review & Editing.

## Acknowledgments

The authors gratefully acknowledge the support from The Japan Foundation, New York (New York, NY) to E.L.D. and to the support from Universidad Católica de Santa María. This was facilitated by the Fondos Internos para la Investigación Tiempo Completo 2025, approved by resolution no. 9313-CU-2025, and through institutional grant nos. 27574-R-2020 and 28048-R-2021 to L.D.G-M. and M.A.C-F.

## Abbreviations

1Abbreviations

DENV: Dengue virus
DENV-NS3: Dengue virus non-structural protein 3
DENV-NS3_hel_: DENV-NS3 helicase/NTPase catalytic domain
DENV-NS3_pro_: DENV-NS3 protease catalytic domain
DEN2-NS3: Dengue virus serotype 2 non-structural protein 3
DEN2-NS3(S171-K618): DEN2-NS3 helicase/NTPase catalytic domain
DMSO: dimethyl sulfoxide
DNase: I deoxyribonuclease I
ECG: (−)-epicatechin gallate
EGC: (−)-epigallocatechin
EGCG: (−)-epigallocatechin gallate
F.A.C.: final assay concentration
HEPES: 4-(2-hydroxyethyl)piperazine-1-ethanesulfonic acid
HTS: high-throughput screening
K_i_: inhibition constant
*Lm*G6PDH: *Leuconostoc mesenteroides* glucose-6-phosphate dehydrogenase
LB: Luria-Bertani
MD: molecular dynamics
MP: mobile phase
NA: nucleic acid
NPI: normalized percent inhibition
NS3: non-structural protein 3
nsp13: non-structural protein 13
NTPase: nucleoside 5’-triphosphatase
PDB: Protein Data Bank
RNase: A ribonuclease A
SARS-CoV-2: Severe Acute Respiratory Syndrome Coronavirus 2
SF1B: Superfamily-1B
SF2: Superfamily-2
TlGlcK: Thermococcus litoralis glucokinase
TEA: triethanolamine
ZIKV: Zika virus
ZIKV-NS3: Zika virus non-structural protein 3

