## Supplemental Information for "Targeting Dengue Virus NS3 Helicase: Biochemical and Computational Evaluation of Catechins from *Camellia sinensis* as Potential Therapeutic Leads^†^"

**Table S1**. DEN2-NS3(S171-K618) – inhibitor $K_i$ values determined from three independent experiments.

| **Inhibitor** | **Replicate 1: $K_i$** | **Replicate 2: $K_i$** | **Replicate 3: $K_i$** | **$K_i$ (Mean ± SD)** |
|---|---|---|---|---|
| EGCG | 450 nM | 450 nM | 300 nM | 400 ± 86.6 nM |
| ECG | 300 nM | 800 nM | 550 nM | 550 ± 250 nM |
| SSYA10-001 | 10.5 μM | 10.0 μM | 10.0 μM | 10.2 ± 0.3 μM |
| EGC | 20.5 μM | 13.5 μM | 21.0 μM | 18.3 ± 4.2 μM |

**Table S2**. SiteMap[a] pocket characterization of DEN2-NS3(S171-K618).

| **Parameter** | **Value** | **Interpretation** |
|---|---|---|
| SiteScore | 1.120 | Favorable pocket quality |
| Dscore | 1.042 | Druggable-like site |
| Size | 212 | Large pocket |
| Volume | 771.750 $Å^3$ | Compatible with large catechins |
| Donor/acceptor | 0.614 | H-bond-compatible environment |
| Enclosure | 0.878 | Moderately enclosed pocket |
| Exposure | 0.489 | Moderate solvent exposure |
| Hydrophilic | 1.303 | Strong polar character |
| Hydrophobic | 0.618 | Lower hydrophobic contribution |

[a] Halgren, T. A. (2009) Identifying and characterizing binding sites and assessing druggability, *J. Chem. Inf. Model. 49*, 377-389.

**Table S3**. Prime MM-GBSA energetic decomposition.[a]

| Compound | $\Delta G_{bind}$ | Coulomb | Covalent | H-bond | Lipo | SolvGB | vdW |
|---|---|---|---|---|---|---|---|
| EGCG | −51.97 ± 0.01 | −28.26 ± 0.86 | +5.43 ± 0.88 | −3.87 ± 0.25 | −16.28 ± 0.97 | +39.76 ± 0.25 | −46.88 ± 0.38 |
| ECG | −53.88 ± 0.52 | −38.56 ± 0.52 | +7.20 ± 0.65 | −3.39 ± 0.67 | −15.76 ± 0.35 | +43.01 ± 0.96 | −45.24 ± 0.82 |
| ATP | +2.98 ± 0.21 | +82.54 ± 0.80 | +5.83 ± 0.65 | −7.37 ± 0.37 | −5.90 ± 075 | −33.50 ± 0.01 | −37.27 ± 0.64 |
| SSYA10-001 | +5.40 ± 0.03 | −12.01 ± 0.96 | +2.98 ± 0.84 | −0.95 ± 0.93 | −10.23 ± 0.63 | +50.11 ± 0.88 | −24.49 ± 0.71 |
| Mosnodenvir | −40.22 ± 0.22 | +22.59 ± 0.84 | +20.13 ± 0.08 | −1.66 ± 0.15 | −18.60 ± 0.33 | −5.01± 0.05 | −55.20 ± 0.67 |

[a] Docking scores and MM-GBSA values are reported as mean ± SD from three repeated docking/rescoring calculations.

**Table S4**. Shared residue interaction map.

| Residue | EGCG | ECG | ATP | Mosnodenvir | SSYA10-001 | Proposed role |
|---|---|---|---|---|---|---|
| ARG599 | yes | yes | yes | water-mediated | yes | recurrent polar anchor |
| ASP603 | yes | strong | yes | weak/absent | weak | catechin/ATP polar node |
| ASP541 | yes | yes | possible | no | no | catechin-associated polar contact |
| ARG387/ LYS388 | yes | yes | yes | weak | yes | charged network |
| ASP409 | yes | transient | yes | strong | yes | recurrent pocket residue |
| HIS487 | yes | yes | yes | yes | strong | shared dynamic contact |
| MET429 | yes | weak | yes | strong | yes | hydrophobic/polar pocket border |
| ASP290 | weak/moderate | strong | possible | no | no | ECG-specific anchor |

**Table S5.** Replica-level residue–ligand interaction occupancies across three independent 200-ns MD simulations.

| Ligand | Residue | Rep 1 (%) | Rep 2 (%) | Rep 3 (%) | Mean ± SD (%) |
|---|---|---|---|---|---|
| EGCG | ARG599 | 62 | 58 | 65 | 61.7 ± 3.5 |
| EGCG | ASP603 | 55 | 51 | 58 | 54.7 ± 3.5 |
| EGCG | ASP541 | 49 | 45 | 50 | 48.0 ± 2.6 |
| EGCG | ARG387 | 47 | 50 | 45 | 47.3 ± 2.5 |
| EGCG | MET429 | 38 | 35 | 40 | 37.7 ± 2.5 |
| ECG | ASP603 | 79 | 83 | 76 | 79.3 ± 3.5 |
| ECG | ASP290 | 67 | 64 | 71 | 67.3 ± 3.5 |
| ECG | ARG599 | 55 | 59 | 52 | 55.3 ± 3.5 |
| ECG | ASP541 | 51 | 47 | 54 | 50.7 ± 3.5 |
| ECG | ARG387 | 46 | 49 | 43 | 46.0 ± 3.0 |
| ECG | VAL544 | 35 | 38 | 33 | 35.3 ± 2.5 |
| ATP | ARG387 | 52 | 48 | 55 | 51.7 ± 3.5 |
| ATP | LYS388 | 47 | 43 | 50 | 46.7 ± 3.5 |
| ATP | ARG599 | 45 | 42 | 47 | 44.7 ± 2.5 |
| ATP | ASP409 | 41 | 46 | 39 | 42.0 ± 3.6 |
| ATP | ASP603 | 39 | 44 | 37 | 40.0 ± 3.6 |
| ATP | HIS487 | 36 | 40 | 34 | 36.7 ± 3.1 |
| Mosnodenvir | ASP409 | 58 | 62 | 54 | 58.0 ± 4.0 |
| Mosnodenvir | MET429 | 49 | 53 | 46 | 49.3 ± 3.5 |

| Ligand | Residue | Rep 1 (%) | Rep 2 (%) | Rep 3 (%) | Mean ± SD (%) |
|---|---|---|---|---|---|
| Mosnodenvir | HIS487 | 37 | 42 | 35 | 38.0 ± 3.6 |
| Mosnodenvir | CYS428 | 33 | 36 | 31 | 33.3 ± 2.5 |
| Mosnodenvir | ILE365 | 29 | 32 | 27 | 29.3 ± 2.5 |
| SSYA10-001 | HIS487 | 57 | 61 | 54 | 57.3 ± 3.5 |
| SSYA10-001 | ARG599 | 43 | 47 | 40 | 43.3 ± 3.5 |
| SSYA10-001 | ASP409 | 39 | 44 | 36 | 39.7 ± 4.0 |
| SSYA10-001 | ARG387 | 35 | 38 | 32 | 35.0 ± 3.0 |
| SSYA10-001 | CYS485 | 31 | 34 | 29 | 31.3 ± 2.5 |

Occupancy was calculated independently for each 200-ns MD replica as the percentage of analyzed trajectory frames in which at least one qualifying residue–ligand interaction was detected. Values are reported as mean ± sample SD across three independent MD replicas (n = 3).

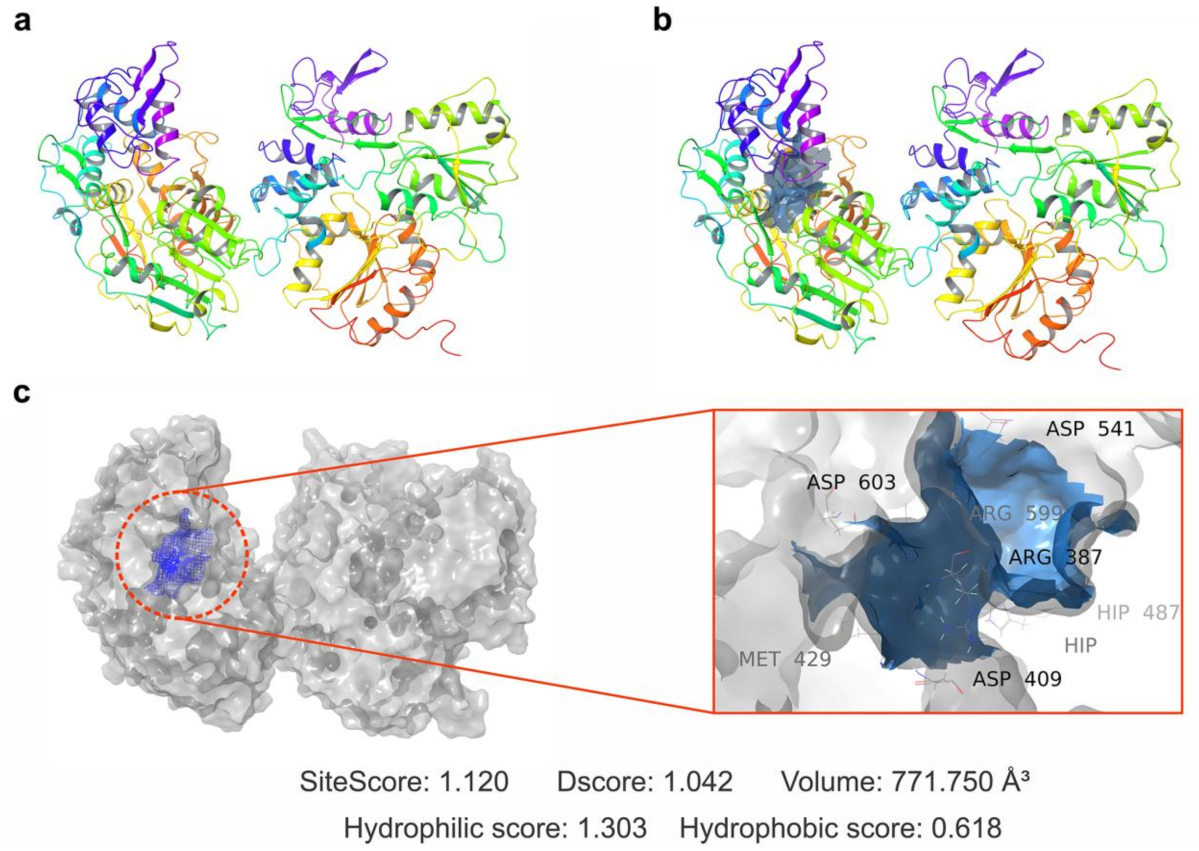


**Figure S1.** A druggable amphipathic cavity composed of residues that form the functional RNA-binding channel in the DEN2-NS3 helicase/NTPase catalytic domain. The principal residues include ASP290, ARG387, ASP409, MET429, HIS487, ASP541, ARG599, and ASP603. (a) Overall structure of DEN2-NS3(S171–K618) based on entry PDB_00002BHR. (b) Surface representation of the pocket used for grid generation and docking. (c) Close-up view of the selected druggable pocket with nearby ligand-recognition residues.

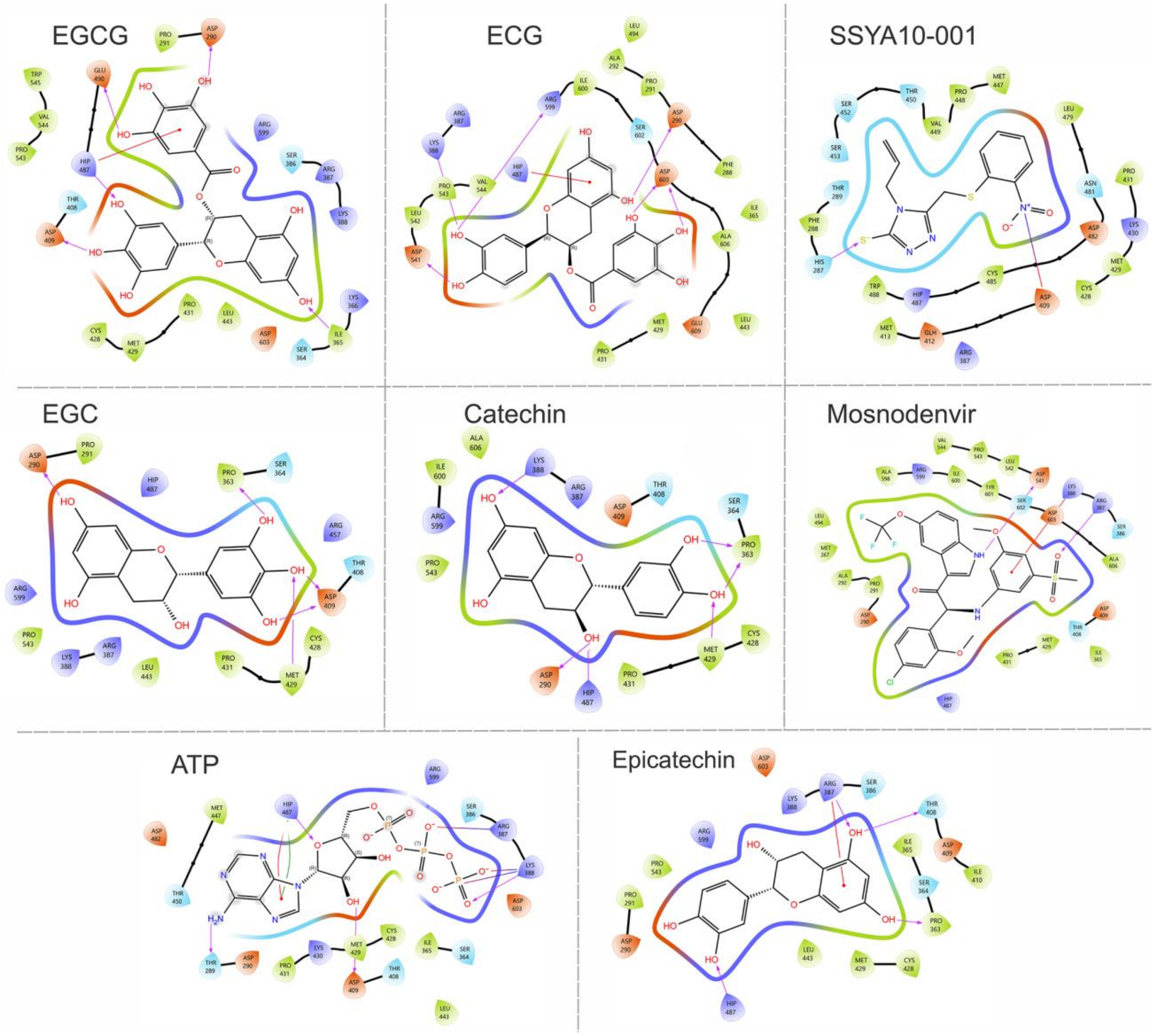


**Figure S2.** Two-dimensional ligand–protein interaction diagrams of catechin derivatives and reference compounds docked into the pocket of DEN2-NS3 helicase/NTPase catalytic domain. Representative 2D interaction diagrams are shown for EGCG, ECG, SSYA10-001, EGC, catechin, mosnodenvir, ATP, and epicatechin within the same pocket.

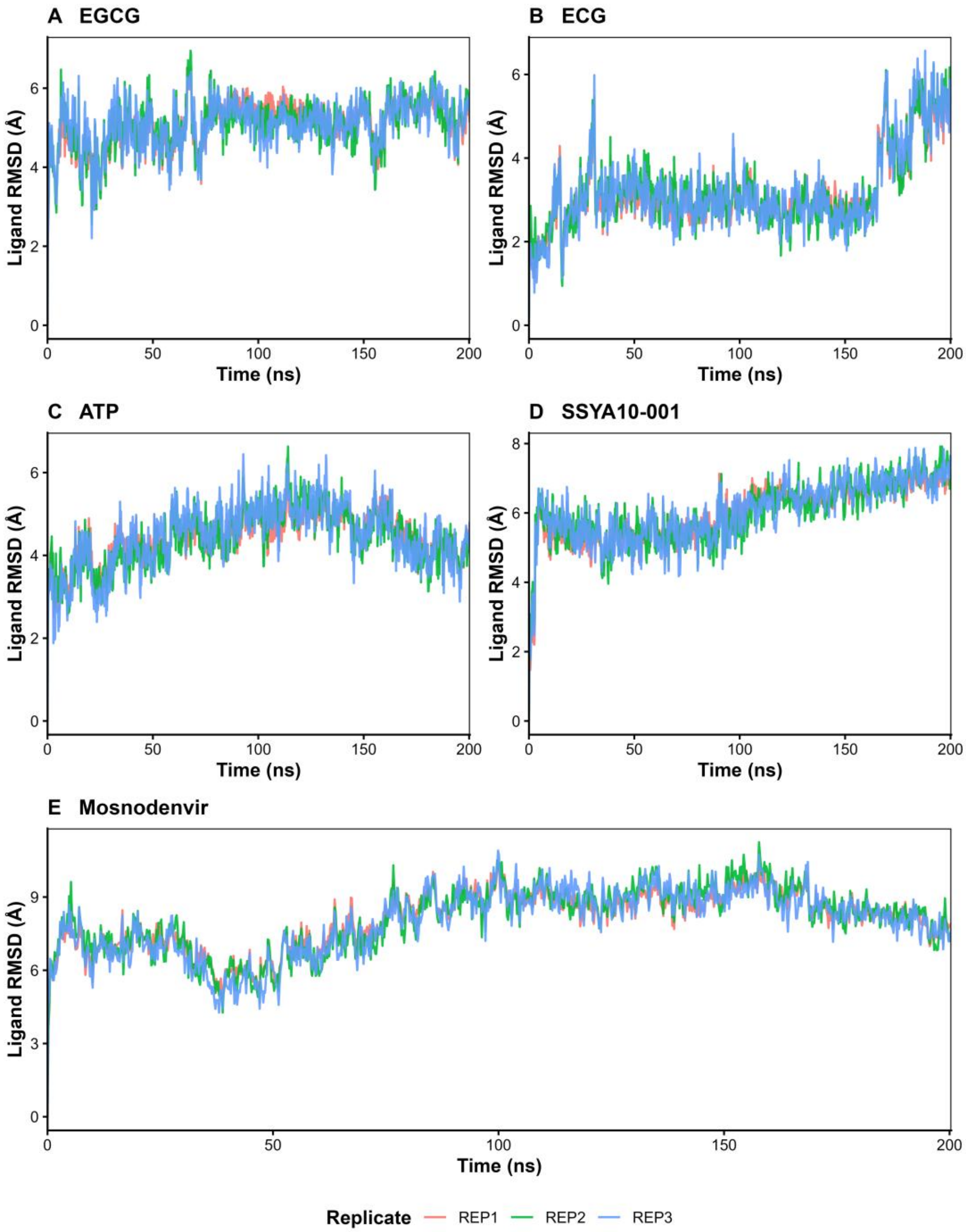


**Figure S3.** Replica-resolved ligand RMSD analysis of selected DEN2-NS3(S171–K618) complexes. Ligand RMSD relative to the protein-aligned frame is shown for three independent 200-ns MD replicas of (A) EGCG, (B) ECG, (C) ATP, (D) SSYA10-001, and (E) mosnodenvir. REP1, REP2, and REP3 represent the individual trajectories used to assess trajectory-to-trajectory reproducibility and conformational variability. These replicate-resolved profiles complement the representative RMSD traces shown in Figure 6.
